# Enhanced inter-regional coupling of neural responses underlies long-term behavioral priming

**DOI:** 10.1101/2020.05.08.084921

**Authors:** Stephen J. Gotts, Shawn C. Milleville, Alex Martin

**Affiliations:** Section on Cognitive Neuropsychology, Laboratory of Brain and Cognition, National Institute of Mental Health, National Institutes of Health, Bethesda, Maryland, 20892, United States of America

**Keywords:** synchrony, predictive coding, implicit memory, functional connectivity, effective connectivity, structural equation modeling, MVPA

## Abstract

Stimulus identification commonly improves with repetition over long delays (“repetition priming”), whereas neural activity commonly decreases (“repetition suppression”). Multiple models have been proposed to explain this brain-behavior relationship, predicting alterations in functional and/or effective connectivity (*Synchrony* and *Predictive Coding* models), in the latency of neural responses (*Facilitation* model), and in the relative similarity of neural representations (*Sharpening* model). Here, we test these predictions with fMRI during overt and covert naming of repeated and novel objects. While we find partial support for predictions of the Facilitation and Sharpening models in the left fusiform gyrus and left frontal cortex, the strongest support was observed for the Synchrony model, with increased coupling between right temporoparietal and anterior cingulate cortex for repeated objects that correlated with priming magnitude across participants. Despite overlap with regions showing repetition suppression, increased coupling and repetition suppression varied independently, establishing that they follow from distinct mechanisms.

**Highlights:**

- Tested four prominent neural models of repetition suppression and long-term priming
- Connectivity analyses supported Synchrony model but not Predictive Coding model
- Timing, spatial similarity of responses partially support Facilitation, Sharpening
- Repetition suppression was independent of coupling, implying distinct mechanisms

**eTOC:** Gotts et al. test four prominent neural models of repetition suppression and behavioral priming. They show that the model with the most support is the Synchrony model: a whole-brain connectivity analysis revealed that temporoparietal cortex has increased coupling with anterior cingulate cortex following repetition, particularly for strongly primed objects.

## Introduction

Repeated exposure to objects during the performance of a task leads to improved identification speed and accuracy, a phenomenon referred to as “repetition priming” (e.g. Cave & Squire, 1992; Scarborough et al., 1977; Tulving & Schacter, 1990). Repetition priming is stimulus-specific, occurs in a wide range of tasks, and is extremely long-lasting, surviving delays of months and even years (e.g., Cave, 1997; Mitchell, 2006; van Turennout, Ellmore, & Martin, 2000). The relative sparing of repetition priming in patients with damage to medial temporal lobe structures, such as those with amnesia, also highlights the likely neocortical basis of the phenomenon (e.g. Graf, Squire, & Mandler, 1984; McClelland, McNaughton, & O’Reilly, 1995; Squire, 1992; Warrington & Weiskrantz, 1974). Indeed, neocortical brain regions commonly exhibit a companion phenomenon to repetition priming referred to as “repetition suppression”, in which neural activity in task-engaged brain regions decreases with repetition (e.g. Desimone, 1996; Henson, 2003; Wiggs & Martin, 1998; Schacter & Buckner, 1998). Like repetition priming, repetition suppression is stimulus-specific, builds up across repetitions, occurs in a wide-range of tasks and can be long-lasting (e.g. Li, Miller, & Desimone, 1993; van Turennout, Bielamowicz, & Martin, 2003; Wiggs, Weisberg, & Martin, 2006). The joint occurrence of repetition priming and repetition suppression across a wide range of experimental contexts with different sensory and motor modalities has led to the notion that these phenomena reflect incremental neocortical learning mechanisms, serving to form and shape long-term perceptual, conceptual, and motor knowledge representations throughout the brain (e.g. Gotts, 2016; McClelland et al., 1995; Newman & Norman, 2010; Norman et al., 2006; Stark & McClelland, 2000).

Multiple theoretical models have been proposed to account for the simultaneous observation of both neural repetition suppression and behavioral priming (reviewed in Grill-Spector, Henson, & Martin, 2006; Gotts, Chow, & Martin, 2012; see Figure 1). The *Synchrony* model (Ghuman et al., 2008; Gilbert et al., 2011; Gotts, 2003; see also Brunet et al., 2014; Engell & McCarthy, 2014) holds that as neural activity decreases, cells become more synchronized in their firing, leading to a larger impact on downstream targets and earlier, more reliable propagation of individual spikes, supporting earlier motor responses. The *Predictive Coding* model (Henson, 2003; Friston, 2005; Friston & Kiebel, 2009) views the cortex as a form of hierarchical generative Bayesian statistical model in which perceptual inference occurs as an interaction between bottom-up sensory input (“evidence”) and top-down expectations (“prediction”). Top-down predictions improve with repetition, reducing prediction error by inhibiting or suppressing bottom-up sensory evidence, thereby producing repetition suppression. Simultaneously, this can speed up evoked neural responses via an increase in synaptic gain due to enhanced encoding precision and confidence, producing behavioral priming. A related model, the *Facilitation* model (Henson, 2003; James et al., 2000; James & Gauthier, 2006), simply posits that evoked neural responses are advanced in time with repetition, accompanied by the earlier termination of activity, reflected as repetition suppression when measured with techniques such as BOLD fMRI. Finally, the *Sharpening* model (Desimone, 1996; Wiggs & Martin, 1998) holds that while neural activity is decreasing overall with repetition, the task-engaged cells are becoming more selectively tuned, with the largest decreases occurring in cells that are poorly responsive and/or weakly tuned to the repeated stimuli. In contrast, cells that are the most responsive and selective to the repeated stimuli maintain their firing rates. When combined, bottom-up support would be removed for alternative or competing representations in downstream brain regions, allowing more rapid propagation of stimulus-selective activity throughout task-engaged neural pathways, as well as faster and more accurate behavioral responses.

**Figure 1.**
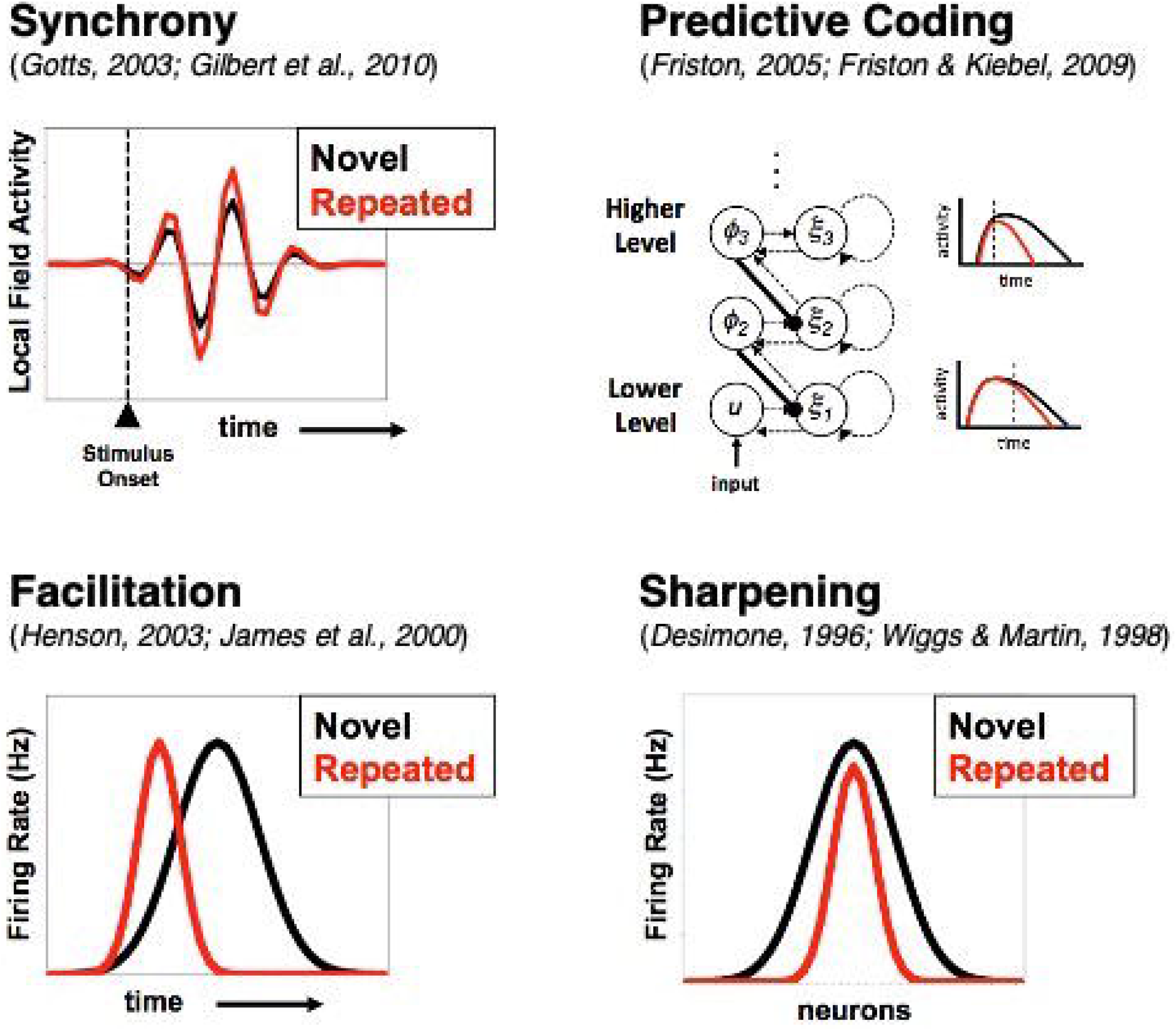
Neural Models of Repetition Priming. Four prominent models of repetition priming and repetition suppression are considered. The Synchrony model (upper left) holds that neural activity becomes more synchronized with repetition, permitting more coordinated propagation of activity at lower overall activity levels. The Predictive Coding model (upper right) holds that top-down causal influences are more strongly negative, leading to repetition suppression in the receiving region, along with a gain enhancement that leads to more rapid onset and offset of responses. The Facilitation model (lower left) claims that neural activity onset and offset are advanced in time, with earlier peak responses and a reduction in overall activity. The Sharpening model (lower right) holds that weakly tuned, poorly responsive cells are the ones driving repetition suppression, reducing downstream support for competing stimulus identities and speeding downstream stimulus-selective responses. Figure reproduced from Gotts, Chow, & Martin (2012) (permission pending).

Common to most of these models is the prediction that measures of connectivity between brain regions should be affected by repetition. The Synchrony model predicts increased positive inter-regional coupling with repetition. The Sharpening model predicts an increase in positive coupling from regions exhibiting repetition suppression, accompanied by a reduction in the similarity of neural responses locally due reduced overlap among neural representations. The Predictive Coding model predicts that top-down connections should have stronger negative coupling with repetition and that this coupling should correlate with the magnitude of repetition suppression in the regions receiving this input. The Facilitation model is less clear regarding predictions about coupling but predicts that neural responses in regions exhibiting repetition suppression should be temporally advanced for repeated objects, with earlier onset and peak. Finally, each model predicts that behavioral priming magnitude should be related to its core changes.

We test each of these predictions in the context of an object identification task in fMRI. Participants were required to name pictured common objects (Figure 2). Overt picture naming has frequently been used in the repetition priming literature (e.g. Bartram, 1973; Cave & Squire, 1992; Gotts et al., 2015; Wiggs et al., 2006; see Francis, 2014, for review), with the advantages that the priming effects are large and correct responses are unlikely to be due to guessing (in contrast to alternative-forced-choice tasks). However, given the risks of motion-related artifacts when employing overt verbal responding, covert naming (e.g. Gilbert et al., 2011; van Turennout et al. 2000), in which participants named objects silently while pressing a response button to mark the naming onset, was also included as a form of artifact control. Features of neural activity that do not differ across overt and covert naming cannot be due to overt speech artifacts. Given the importance of between-region connectivity to the predictions being tested, a novel form of fMRI task-based connectivity was designed to eliminate contamination by the temporal contour of the task-evoked response to functional and effective connectivity measures. A slow event-related design was used, permitting relatively clean isolation of the peak response to each individual trial compared to rapid event-related designs. The response peak was then notched out of the time series, with connectivity calculated as the co-fluctuation (correlation) of peak responses across trials between pairs of regions/voxels.

**Figure 2.**
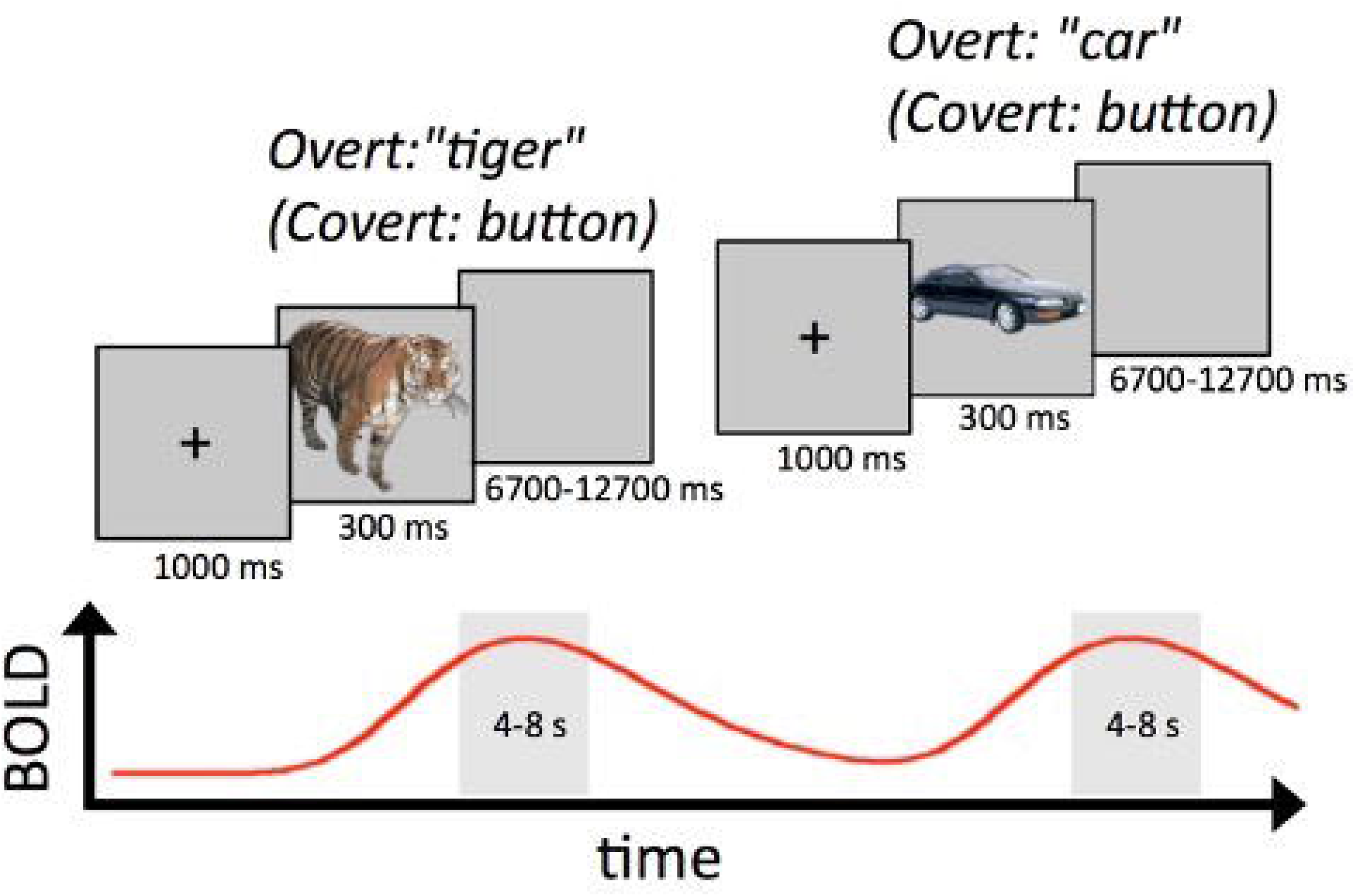
Overt and Covert Picture Naming Tasks. Participants were instructed to name pictures out loud (Overt Naming) or silently to themselves, pressing a button to mark the naming time (Covert Naming). Trials were structured in a slow event-related design in order to separate the peak responses of individual trials, and jitter of 8 to 14 seconds occurred from trial onset to trial onset. Name responses were marked for correctness and the onset time of the voice/button response was recorded for each trial. For analyses of local activity, an empirical hemodynamic response function was estimated across trials with a separate regressor at each time point following stimulus onset. For connectivity analyses, the peak BOLD response was calculated for each trial and in each fMRI voxel by averaging the 2 timepoints (4-6 and 6-8 seconds) adjacent to the expected peak of the hemodynamic response function, with a single peak value saved for each trial in order to eliminate the contribution of the temporal contour of the evoked responses from functional and effective connectivity estimates.

## Results

### Regions Engaged in Object Naming and Showing Repetition Effects

Task analyses first identified brain regions commonly engaged in overt and covert naming (32 and 28 participants, respectively; see STAR Methods). Voxels with above-baseline responses in both tasks are shown in Figure 3A at two levels of significance, one with a minimum level of significance (P<.05, FDR-corrected to q<.05; shown in orange) and one with a more stringent level of significance in both tasks individually, for which responses can be said to replicate across tasks (P<.0001, q<.00016 in each task; shown in red). As in previous studies of picture naming responses (e.g. Gilmore et al., 2019; Kan & Thompson-Schill, 2004; van Turrenout et al., 2000, 2003), these regions included left and right lateral prefrontal cortex, bilateral occipitotemporal and ventral temporal cortex, bilateral intraparietal cortex, anterior cingulate cortex, thalamus, striatum, and cerebellum.

**Figure 3.**
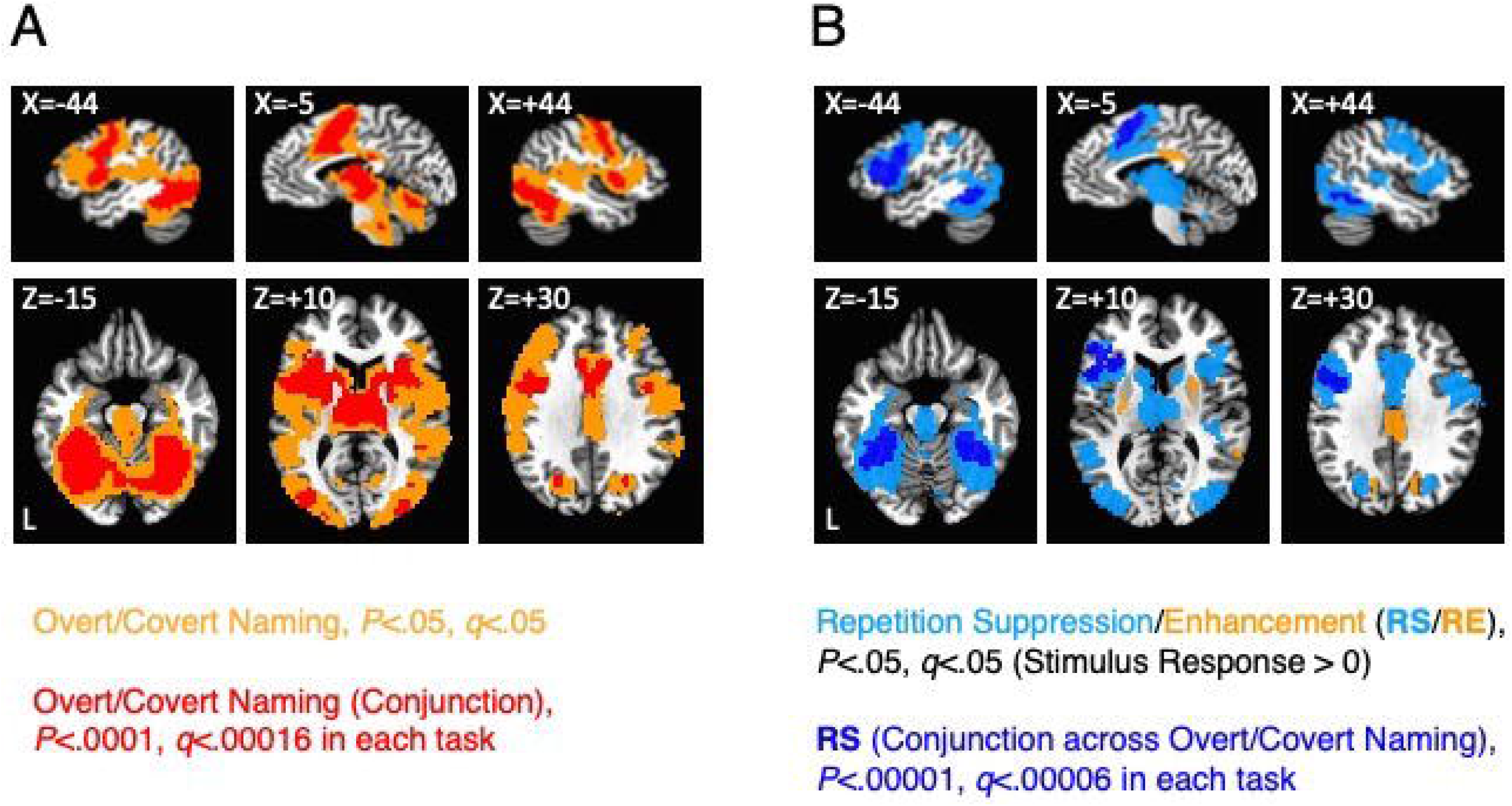
Regions Engaged in Object Naming and Showing Repetition Effects. (A) Above-baseline responses to Overt and Covert Naming tasks are shown at two levels of significance, one at a minimum level of significance (pooling the tasks, P<.05, FDR q<.05; shown in orange) and another at a more restrictive level of significance (P<.0001, FDR q<.00016 in each task individually) for which responses can be said to replicate across tasks (shown in red). (B) Repetition effects, either repetition suppression (blue colors) or repetition enhancement (orange), are shown at two levels of significance, one at a minimum level of significance (pooling the tasks, P<.05, FDR q<.05; suppression shown in light blue, enhancement shown in orange) and another at a more restrictive level of significance (P<.00001, FDR q<.00006 in each task individually). Repetition effects were masked by the less restrictive threshold in (A), as all theories being evaluated make claims about regions engaged at above-baseline levels during the tasks.

Repetition effects were then calculated in all voxels showing above-baseline task responses, as all models being tested posit that the relevant changes occur among cell populations engaged by the task. Regions exhibiting repetition effects, either repetition enhancement or repetition suppression, are shown in Figure 3B at two levels of significance. At a minimum level of significance (P<.05, q<.05), most of the voxels responding above baseline exhibited repetition suppression (light blue), with more restricted locations showing repetition enhancement, including cingulate cortex just dorsal to the posterior portion of the corpus callosum, bilateral cuneus, right posterior superior temporal gyrus, and bilateral putamen. At a more stringent level of significance (P<.00001, q<.00006, in each task individually), only repetition suppression was observed across both tasks in four large regions (dark blue): left frontal, bilateral fusiform, and anterior cingulate cortex. Given the concordance of responses in these regions in both overt and covert naming, these findings cannot be easily attributed to overt speech artifacts that would only be present in overt naming.

### Repetition Priming and Primeability in Object Naming

In order to examine correspondence of fMRI responses to behavioral priming magnitude, a within-participant measure of repetition priming magnitude was designed. Examination of the single-participant response times to each object across the three pre-fMRI presentations revealed highly variable responses that only became more reliable when either averaging across participants (Figure 4A and B) or when pooling objects within-participant into larger groupings (Figure 4C; see SI, Figure S1). This fact, combined with the need to pool trials into larger groups anyway in order to calculate connectivity estimates across trials (with only a single datapoint contributed by each trial), led us to adopt a median split of trials into strongly and weakly primed objects. The selection of objects into strongly and weakly primed groupings was actually based on the group-average naming time to an object when NEW, as this variable was strongly and directly related to eventual priming magnitude (Fig. 4B). Since this distinction is defined on normative group measurements rather than on an individual participant’s measured priming, we refer to it as “primeability” rather than priming, in analogy to recent studies of “memorability” as an item property for explicit recollection (e.g. Bainbridge, 2017; 2020; Bainbridge & Rissman, 2018). Observed priming magnitude for each participant on strongly primeable and weakly primeable objects was then quantified by effect size (Cohen’s *d*) in order to adjust for differences in raw response time per participant and per condition. As expected based on the object grouping scheme, strongly primeable objects exhibited larger priming effect sizes than weakly primeable objects, both for overt naming times during fMRI and button-press times for covert naming during fMRI (Fig. 4C).

**Figure 4.**
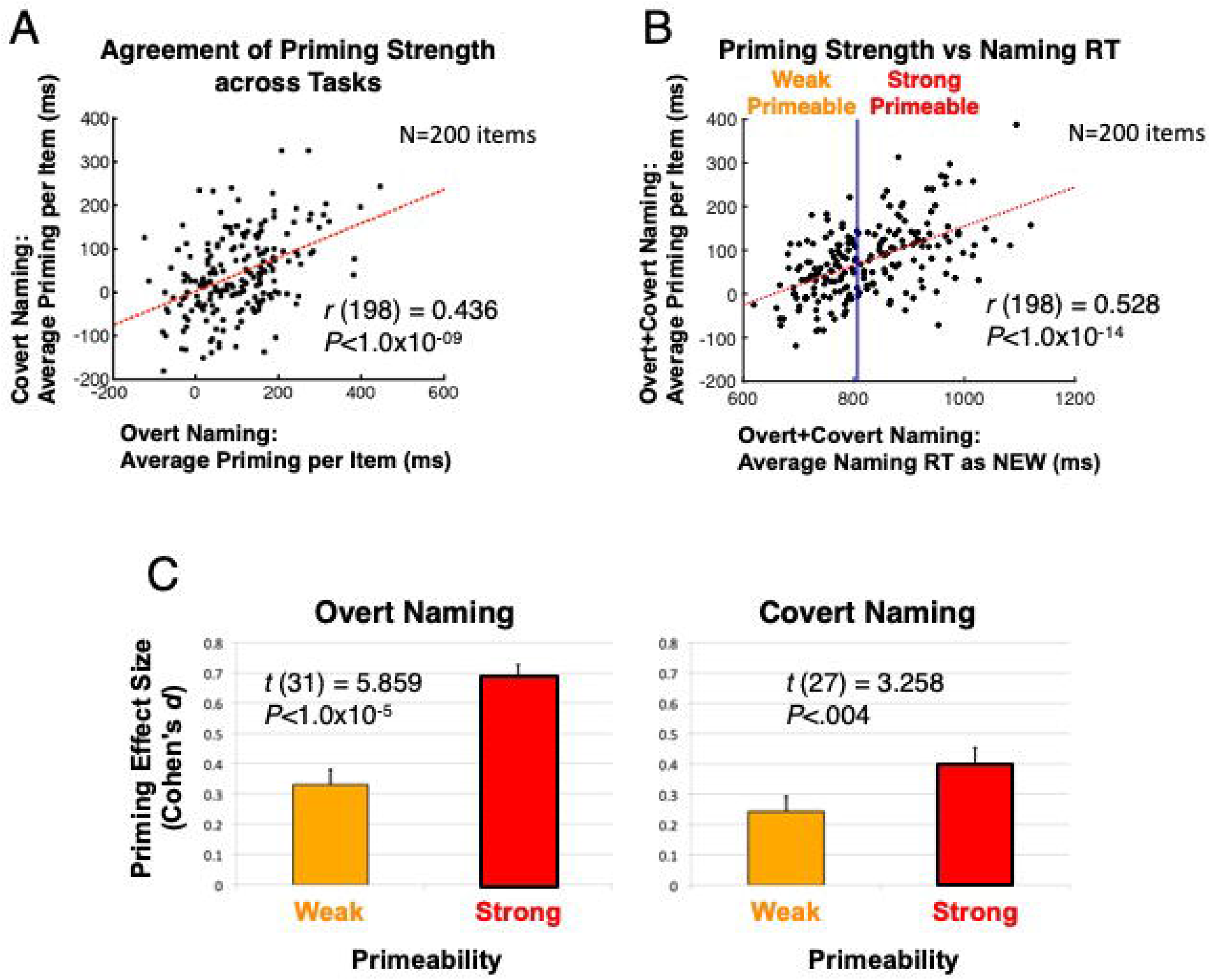
Repetition Priming and Primeability. (A) When averaged across participants within each task, priming magnitudes specific to each of the 200 stimuli are highly reliable across task. For the covert naming task, these priming magnitudes were assessed in a post-fMRI session during which participants overtly named all stimuli (either pre-exposed as OLD or novel to the fMRI session, NEW). (B) When responses are averaged across both participants and tasks, a strong relationship was observed between average response time (RT) to an object when NEW and the subsequent priming magnitude observed. Items were subjected to a median split based on the RT when NEW (using the group-average normative data) to classify objects as either Strong Primeable (slow RT) or Weak Primeable (fast RT), permitting a within-participant measure of “Primeability”. (C) When grouping objects by Primeability (either Strong or Weak), priming effect sizes for each participant were indeed greater for Strong compared to Weak Primeable objects in the Overt Naming task (left panel); this is expected since these responses contributed to the original calculation of Primeability. However, this same relationship was also found for the Covert Naming button press response times during fMRI, which were not used in calculating Primeability (right panel). For related content in SI, see Figure S1.

### Functional Connectivity Changes Related to Primeability

Rather than restricting the evaluation of functional connectivity changes to those regions exhibiting repetition suppression or enhancement, we instead performed a whole-brain search at the voxel level using “connectedness” (e.g. Gotts, Simmons, Milbury, et al., 2012; Jasmin et al., 2019; Berman et al., 2016; Song et al., 2015; see also Cole et al., 2010; Salomon et al., 2011). This approach simplifies the bivariate map of correlations among all possible pairs of voxels into a univariate map of average correlations, with each voxel’s value representing the average functional connectivity level with a desired cohort of voxels, in this case the set of all task-responsive voxels (see STAR Methods). Effects in whole-brain connectedness can then identify effective seeds for more traditional seed-based analyses, thereby detecting a more complete set of regions exhibiting effects in functional connectivity (e.g. Gotts, Simmons, Milbury, et al., 2012; Jasmin et al., 2019; Berman et al., 2016; Song et al., 2015; Steel et al., 2016; Stoddard et al., 2016; Watson et al., 2019).

Voxelwise connectedness estimates were calculated for each participant in four conditions: Strong Primeable OLD, Strong Primeable NEW, Weak Primeable OLD, and Weak Primeable NEW (e.g. a Strong Primeable OLD object corresponded to an object named pre-fMRI and that was strongly primeable, i.e. had a slower than median response time in the normative picture naming data; a Strong Primeable NEW object corresponded to an object that was strongly primeable based on the normative data, but had not been previously seen). These voxelwise estimates were then entered into a linear mixed effects model with factors of Task (Overt, Covert), Primeability (Strong, Weak), and Repetition (OLD, NEW), covarying the global correlation level, GCOR (Saad et al., 2013; Gotts et al., 2013), per condition and participant as a measure of residual whole-brain artifacts (e.g. Gotts et al., 2020; Jasmin et al., 2019; Saad et al., 2013; Zachariou et al., 2017). There was no overall main effect of Repetition that survived whole-brain correction, but there was a significant interaction between Repetition and Primeability in the right temporoparietal cortex (P<.001, corrected to P<.025; Figure 5A). The 3-way interaction with Task failed to reach significance in this or any other location, which is to say that the interaction between Repetition and Primeability was not found to differ significantly in overt versus covert naming. Using this temporoparietal region as a seed and testing a corresponding seed-based linear mixed effects model, significant Repetition X Primeability interactions (P<.001, corrected to P<.025) were observed with the right fusiform gyrus, the anterior cingulate, right STG, and right putamen. To these regions, we added all regions exhibiting significant repetition effects locally that replicated across tasks, given their importance to all theories being tested (Figure 5B). This led to a total of 7 regions of interest (ROIs), sampled as spheres centered on the peak statistic from each region: 3 ROIs identified as showing effects in functional connectivity (right temporoparietal, right STG, and right putamen), 2 ROIs showing repetition suppression effects (left frontal cortex and left fusiform gyrus), and 2 ROIs showing both functional connectivity and repetition suppression effects (ACC and right fusiform gyrus).

**Figure 5.**
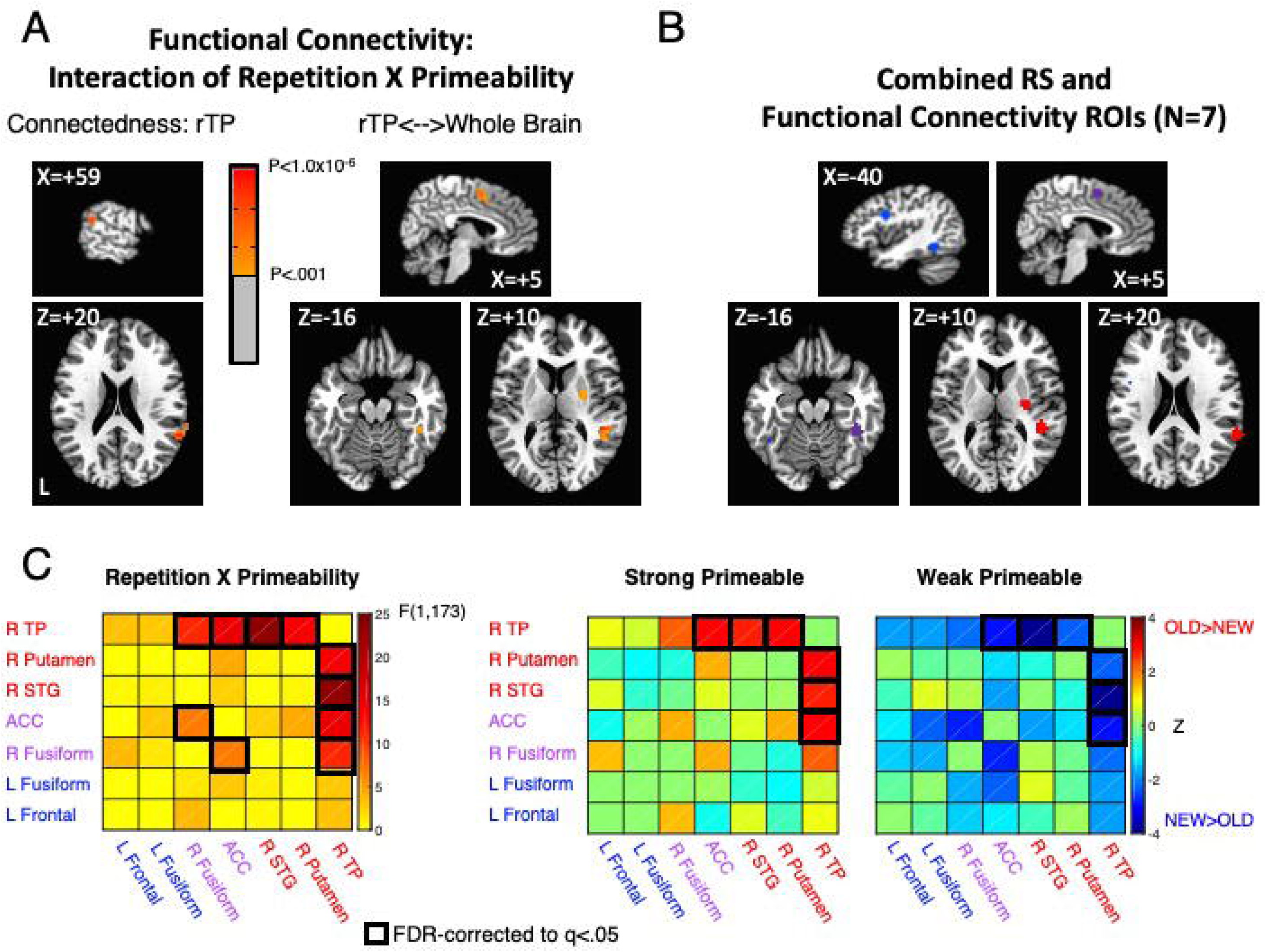
Functional Connectivity Shows Interaction between Primeability and Repetition. (A) A right temporoparietal region (rTP) exhibited an interaction in whole-brain connectedness between Repetition and Primeability, similar to that seen in the behavioral priming results (P<.001, corrected to P<.025). When used as a seed, rTP jointly exhibited this interaction with the anterior cingulate (ACC), right putamen, right STG, and the right fusiform gyrus. (B) Regions showing a Repetition X Primeability interaction were combined with regions showing repetition suppression in both Overt and Covert Naming Tasks. (C) Region-by-region functional connectivity interactions of Repetition X Primeability are shown for all 7 regions (left panel). FDR-corrected effects are indicated by black boxes (P<.01, q<.05). Region-by-region comparisons of OLD versus NEW functional connectivity are shown for Strong and Weak Primeable conditions separately in the right panels. Warm colors (red) indicate OLD > NEW and cool colors (blue) indicate NEW > OLD. FDR-corrected effects (P<.0066, q<.05) were calculated among all region-by-region combinations showing a significant Repetition X Primeability interaction, indicated by black squares. For related content in SI, see Figure S2 and Table S1.

The ROI-level data were submitted to an additional linear mixed effects analysis, affording the assessment of all region-by-region effects of repetition on functional connectivity, as well as interactions between Task, Primeability, and Repetition. As with the voxelwise data, there was no overall effect of Repetition among the ROIs. There was a significant interaction of Repetition and Primeability involving the same regions as detected in the whole-brain analysis (as expected), but there was also an additional interaction detected between the ACC and right fusiform gyrus (all P<.0099, q<.05; Figure 5C). When evaluating the qualitative nature of these interactions by separately assessing Repetition in the Strong Primeable and Weak Primeable conditions, greater functional connectivity was observed for OLD compared to NEW objects in the Strong Primeable condition (right temporoparietal ROI with ACC, right putamen, and right STG ROIs), whereas weaker functional connectivity was observed for OLD compared to NEW objects in the Weak Primeable condition (also right temporoparietal ROIs with ACC, right putamen, and right STG ROIs) (all P<.0065, q<.05). As with the whole-brain data, none of these effects exhibited a further interaction with Task (overt versus covert naming).

### Effective Connectivity Changes Related to Primeability and Repetition Suppression

The functional connectivity effects were then further probed with estimates of effective connectivity among the 7 ROIs. This is important because functional connectivity measures are ambiguous with respect to claims about underlying coupling between regions, as localized real changes in coupling can manifest indirectly at other locations through polysynaptic interactions (see Friston, 2011; Reid et al., 2019, for discussion). We therefore applied structural equation modeling (SEM), a form of effective connectivity estimation that utilizes the pattern of correlation among ROIs in order to evaluate directional changes in underlying inter-regional coupling (e.g. Chen et al., 2011; McIntosh & Gonzalez-Lima, 1994; Price et al., 2009). We first performed a search for the optimal SEM model while pooling all data conditions (Task, Primeability, Repetition) (see STAR methods). The optimal model was then parameterized for each participant’s data per condition, with model parameters tested using the same linear mixed effects modeling approach as for tests of functional connectivity. The optimal model is shown in Figure 6A, with arrows indicating non-zero parameters and the direction of the causal interactions among the ROIs. In all, 10 parameters were found to differ significantly from zero, with all connections being positive (above zero) and with no connections showing negative parameters that would be consistent with inhibition or some form of suppression in any of the conditions. Connections exhibiting a significant Repetition X Primeability interaction are shown as thick red arrows (P<.0063, q<.05 for all), with non-significant interactions shown in black. Figure 6B clarifies the nature of these interactions. The connection from the right temporoparietal ROI (R TP) to the ACC ROI exhibited a significantly larger parameter for OLD than NEW objects for the Strong Primeable condition (P<.0035, q<.05), with no significant difference observed for the Weak Primeable condition. In contrast, the connections from the ACC to the right fusiform ROI and the right STG to the R TP ROI exhibited no significant OLD/NEW effect for the Strong Primeable condition and significantly smaller parameter values for OLD compared to NEW objects in the Weak Primeable condition (P<.0091, q<.05 for both). Of these connections, the one from the R TP to the ACC ROI appeared to provide the best candidate for a correlate of priming magnitude, with the strongest effects observed in the Strong Primeable condition (compare parameter values in Figure 6C to priming measures in Figure 4C). We therefore examined the relationship between the inter-participant variability in priming effect sizes and the R TP to ACC ROI parameter values. There was a significant correlation between the contrast in Strong versus Weak Primeable effect sizes in priming (behavioral data) and the SEM parameter values (fMRI data) across participants [Pearson’s *r* (56) = 0.3303, P<.0114]. This correlation appeared to be driven by the data in the Strong Primeable condition [Pearson’s *r* (57) = 0.2692, P<.04; for Weak Primeable condition: *r* (57) = 0.0995, P > .45].

**Figure 6.**
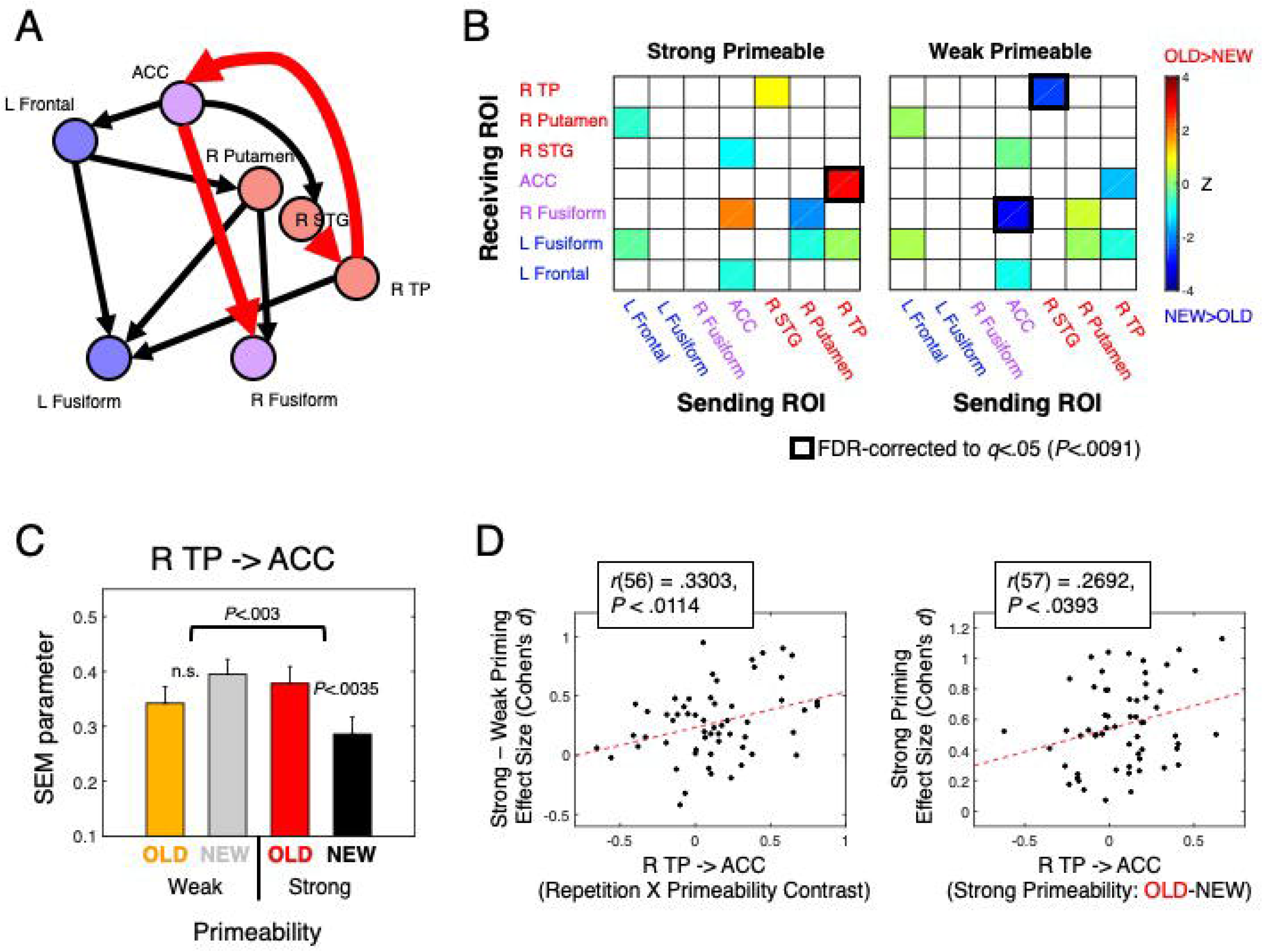
Effective Connectivity Increases between Right Temporoparietal Cortex and ACC correlate with Priming Magnitude. (A) Structural Equation Modeling (SEM) was used to estimate effective connectivity among the 7 Regions. The optimal 10-parameter SEM model is shown, with arrows indicating causal directionality and connections exhibiting a significant Repetition X Primeability interaction (P<.0063, q<.05) shown with thick red arrows (non-significant interactions shown with black arrows). (B) Region-by-region comparisons of SEM parameters for OLD versus NEW objects are shown separately for the Strong and Weak Primeable conditions. For directionality, sending ROIs are listed along the x-axes and receiving ROIs are listed along the y-axes. FDR-corrected OLD/NEW comparisons (P<.0091, q<.05) were assessed among connections exhibiting a Repetition X Primeability interaction, indicated with black squares. (C) The connection from the R TP ROI to ACC ROI shows a Repetition X Primeability interaction that is consistent with the Synchrony model (with increased coupling for OLD objects in the Strong Primeable condition). (D) The R TP to ACC connection further exhibited a correlation across participants with observed priming magnitude, assessed by effect size (Cohen’s *d*). This correlation appeared to be driven by the Strong Primeable condition (rightmost panel), with priming effect size in the Strong Primeable condition correlated with the difference between OLD and NEW SEM parameters in the Strong Primeable condition. For related content in SI, see Figure S3.

These data are most compatible with the Synchrony model, for which coupling is predicted to be greater (and positive) for the OLD compared to the NEW condition, particularly for Strongly Primeable objects. The increased coupling for the OLD condition also occurred despite no change in activity levels in the R TP ROI and significantly reduced activity in the ACC ROI (see SI, Figure S3). The data are less compatible with the predictions of the Predictive Coding model, in which top-down connections are claimed to be more strongly negative following repetition, leading to repetition suppression in the receiving area. As noted, none of the model connections in any of the conditions were found to be negative. Nevertheless, two of the ROIs exhibiting significant repetition suppression (ACC and right fusiform gyrus) were found to have incoming SEM connections modulated by stimulus repetition (from R TP and ACC ROIs, respectively). We therefore examined the relationship between SEM parameter values at these connections and the magnitude of repetition suppression observed in the receiving ROIs. A contrast of repetition suppression in Strong minus Weak Primeable conditions failed to correlate with the corresponding contrast of the SEM parameters (OLD – NEW parameter values in the Strong Primeable condition minus the same difference in the Weak Primeable condition), either for the connection from R TP to ACC [*r* (56) = 0.0897, P>.5] or for the connection from ACC to the right fusiform gyrus [*r* (56) = −0.0429, P>.74]. These results, taken together with the lack of modulation by stimulus repetition of connections into other ROIs exhibiting repetition suppression (e.g. left frontal and left fusiform ROIs), suggest that changes in coupling due to repetition are largely independent of local repetition suppression magnitudes. In order to establish this relationship more clearly for the one connection correlated with behavioral priming magnitude, R TP to ACC, we used partial correlation to remove the magnitude of repetition suppression exhibited in the ACC from the correlation between priming effect size and SEM parameter contrasts in the Strong versus Weak Primeable conditions. This partial correlation remained at approximately the same level as without the partialling [partial *r* (55) = 0.334, P<.0112], establishing that changes in coupling between R TP and ACC are correlated with priming magnitude in a manner largely independent of the magnitude of repetition suppression.

### Testing Additional Predictions of the Facilitation and Sharpening Models

In the previous analyses, we found that functional and effective connectivity effects due to repetition provided critical support for the Synchrony model and failed to provide support for the Predictive Coding model. One aspect of these results also failed to support the Sharpening model, namely that regions showing repetition suppression (L Frontal, L Fusiform, R Fusiform, and ACC ROIs) failed to exhibit increased feed-forward coupling during the processing of OLD relative to NEW objects. Nevertheless, there remain two sets of predictions related to the Facilitation and Sharpening models that can be assessed using the current data.

The Facilitation model predicts that the timing of neural activity, particularly the onset and timing of the peak response, should track repetition suppression and behavioral priming magnitudes. To evaluate this, we extracted the beta coefficients for each participant at each time point following stimulus onset in each of the ROIs exhibiting repetition suppression. A continuous response function was fit to these data points in each of the four conditions (OLD, NEW X Strong, Weak Primeable), affording separate estimates of the timing and amplitude of the peak BOLD response (see STAR methods). Figure 7 shows the estimated hemodynamic response functions for each of the 4 repetition suppression ROIs, along with estimates of the timing of the peak responses (shown as vertical dotted lines). By definition, the peak amplitudes in the OLD condition are smaller than those in the NEW condition (P < 2.1×10^−05^ for all), although the selection of the ROIs provided no bias as to the differential magnitudes in the Strong versus Weak Primeable conditions, nor was there intrinsic bias in the timing of the peak responses in any of the conditions. The Repetition X Primeability interaction in the peak amplitude of the BOLD response was highly significant in the Left Fusiform ROI [F(1,174) = 10.898, P < .0013, FDR q < .05], with stronger repetition suppression in the Strong (Z = 13.419, P < 1.0×10^−15^) relative to the Weak Primeable condition (Z = 7.185, P < 1.0×10^−12^) and no further interaction with Task (Overt/Covert) (P > 0.2). There was no significant Repetition X Primeability interaction in the timing of the peak responses in any of the 4 ROIs (P > 0.27 for all), but there was a significant main effect of Repetition on the timing of the peak in the Left Fusiform ROI [mean (SE) OLD *t*_*peak*_ = 4.382 (0.0967) sec; NEW *t*_*peak*_ = 5.004 (0.091) sec; F(1,174) = 9.233, P < .0028, FDR q < .05] and an uncorrected effect of Repetition in the Left Frontal ROI [OLD *t*_*peak*_ = 4.647 (0.168) sec; NEW *t*_*peak*_ = 4.979 (0.1257) sec; F(1,174) = 4.849, P < .03, FDR q > .05]. Given the significant Repetition X Primeability interaction in the peak amplitude of the Left Fusiform ROI that was qualitatively similar to the difference in behavioral priming effect sizes, we examined the correlation of repetition suppression magnitude (Strong minus Weak Primeable conditions) and behavioral priming magnitude (effect size in Strong minus Weak Primeable conditions) across the 60 participants. This correlation failed to reach significance [*r* (58) = 0.123, P > 0.35]. Taken together, the earlier peak timing of OLD relative to NEW trials in the Left Fusiform ROI (with a similar uncorrected effect in the Left Frontal ROI) is consistent with predictions of the Facilitation model. However, there was no evidence of earlier response onset (as opposed to peak) on OLD trials and no evidence of association with priming magnitudes by condition or across participants in either peak amplitude or peak timing, providing only partial support for this model.

**Figure 7.**
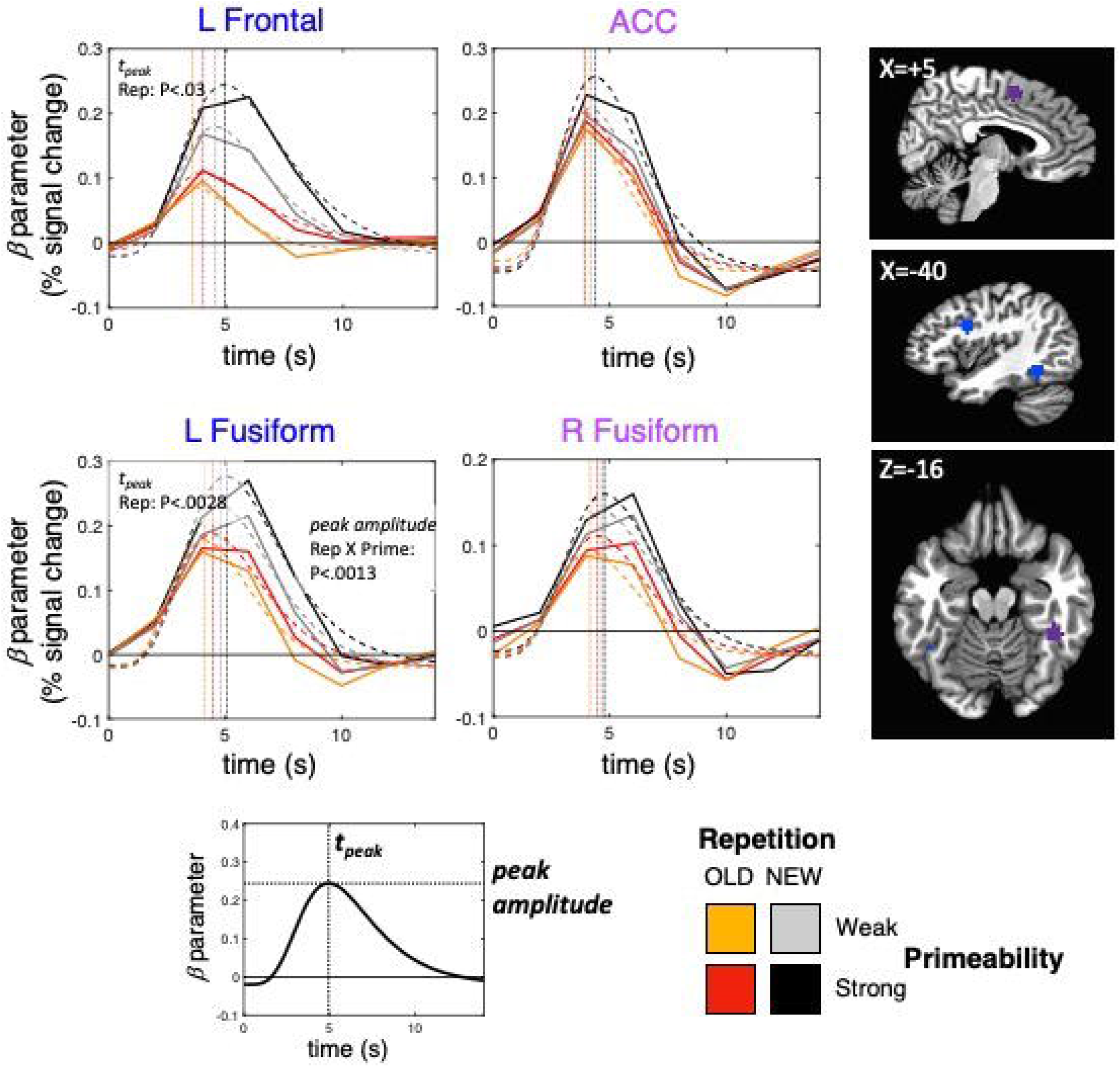
Assessing Activity Timing Predictions of the Facilitation Model. A hemodynamic response function model (gamma variate) was fit to the beta coefficients at each timepoint (TR) for each participant and experimental condition, permitting separate estimates of the peak amplitude and peak time (*t*_*peak*_) (see graphic at bottom). Group average response functions are shown with dashed lines for each condition, and group average estimates of peak times are shown with vertical dotted lines (mean data on the actual measured beta coefficients are shown with solid lines). Main effects of Repetition on peak time were observed in the Left Fusiform and Left Frontal ROIs (OLD peaks earlier than NEW peaks), but there were no significant interactions between Repetition and Primeability on peak time. A significant interaction of Repetition X Primeability was observed in the Left Fusiform ROI for peak amplitude, but this failed to relate to priming magnitude across participants. A color key for the experimental conditions is shown at the bottom right (OLD, Strong Primeable = red; OLD, Weak Primeable = orange; NEW, Strong Primeable = gray; NEW, Weak Primeable = black).

The Sharpening model predicts that the spatial similarity of neural responses should decrease with repetition as activity to OLD stimuli becomes “sharper” and overlaps less across objects. We evaluated this prediction using multi-voxel pattern analysis (MVPA; e.g. Haxby et al., 2001; Misaki et al., 2010; Norman et al., 2006; Op de Beeck et al., 2008) applied to the 4 repetition suppression ROIs. The pattern of peak BOLD responses across voxels in an ROI to each trial was correlated (Pearson) with all correct trials of the same type (e.g. Strong Primeable OLD trials) (see STAR methods). As shown in Figure 8A, the inter-item correlation level was indeed lower for OLD compared to NEW trials (Z > 4.352, P < 1.4×10^−5^ for all 4 ROIs) with a significantly larger decrease for Strong compared to Weak Primeable conditions in the Left Frontal [Repetition X Primeability F(1,174) = 23.259, P < 3.1×10^−6^, FDR q < .05] and ACC ROIs [Repetition X Primeability F(1,174) = 5.530, P < .0199, FDR q < .05]. However, on further examination, these effects appeared to be related to average beta weight levels and estimates of the signal-to-noise ratio (SNR) of the peak responses (see SI, Figure S4). When covarying these quantities, only the Repetition X Primeability interaction in the Left Frontal ROI remained significant [F(1,172) = 16.906, P < 6.1×10^−5^, FDR q < .05], driven by an OLD/NEW difference in the Strong Primeable condition [Z = 3.607, P<;3.2×10^−4^ vs Weak: Z = −.649, P>.5] (Figure 8B). However, even this last effect remains in doubt, given the strong dependence on beta weight levels and SNR for this ROI [β: F(1,172) = 16.117, P < 8.9×10^−5^; SNR: F(1,172) = 19.234, P < 2.1×10^−5^]. In other words, the unadjusted MVPA results are consistent with the predictions of the Sharpening model, but the results may simply reflect the average amplitude of the BOLD response rather than the pattern of responses across voxels, per se, with response levels nearer to zero being more contaminated with noise (see also Ramirez, 2018; Tong et al., 2012, for discussion). Consistent with this interpretation, there was no correlation between MVPA measures and behavioral priming magnitudes across participants in the Left Frontal and ACC ROIs either prior to or after adjustment (| *r* | < 0.1, P > .56 for all).

**Figure 8.**
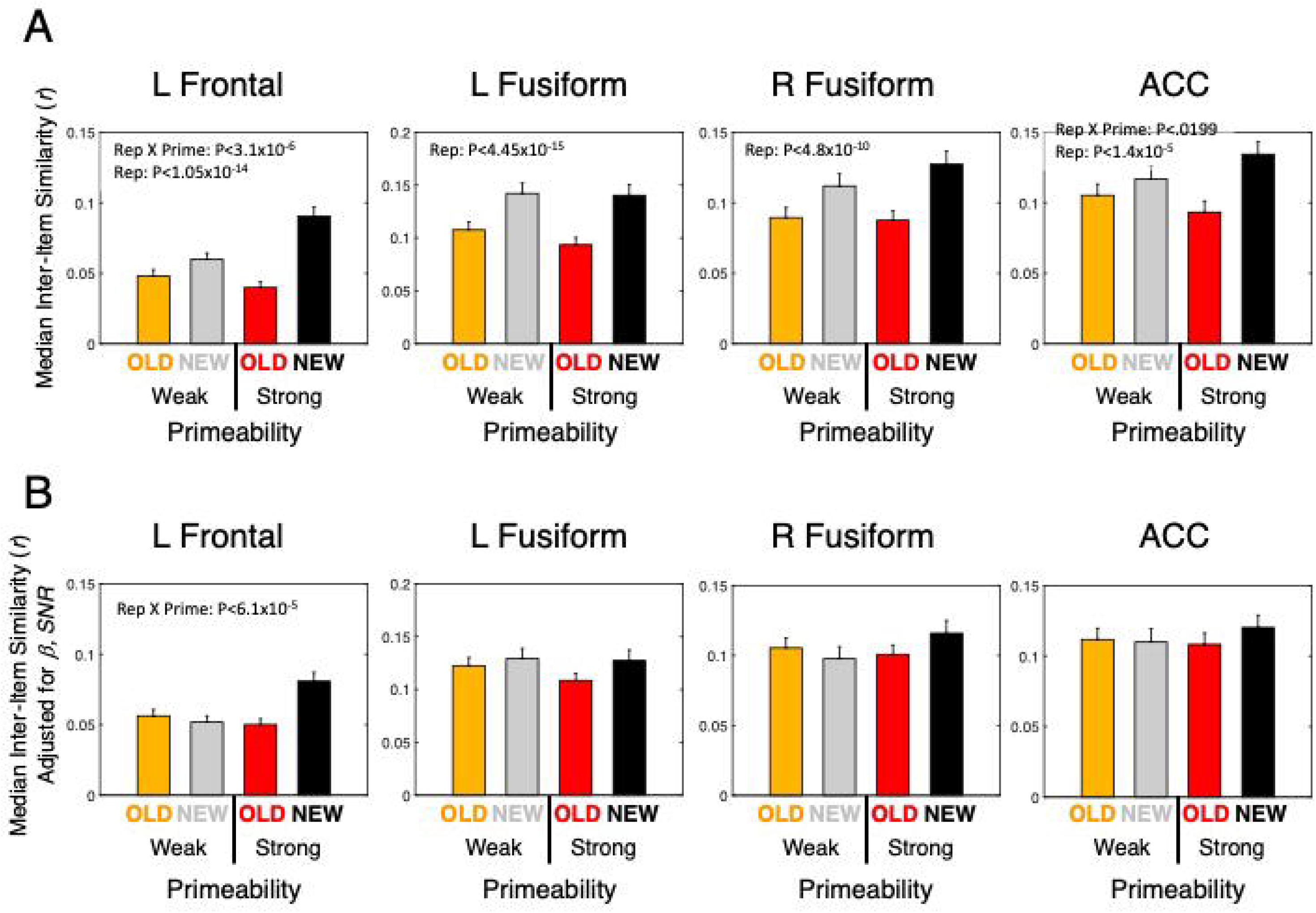
MVPA Tests of the Sharpening Model. (A) Spatial correlations of peak responses on each individual trial were calculated across all trials per condition type using the full extent of the four RS clusters (see Table S1), retaining the median inter-item correlation per participant. A significant Repetition X Primeability interaction was observed for the Left Frontal and ACC ROIs (corrected by FDR, q<.05). (B) Strong covariation of the spatial correlations with average beta coefficients and estimated signal-to-noise ratio eliminated most of the differences between conditions after adjustment for these variables, leaving only a significant Repetition X Primeability interaction in the Left Frontal ROI. For related content in SI, see Figure S4.

## Discussion

In the current study, we have tested predictions from four prominent models (Synchrony, Predictive Coding, Facilitation, and Sharpening) of the joint relationship between repetition suppression and long-term behavioral priming effects. We performed a whole-brain analysis of changes in functional and effective connectivity related to a within-participant measure of priming magnitude referred to as “primeability”, as these changes were central to multiple models. We then tested predictions about the timing of neural responses derived from the Facilitation model, as well as predictions about the decreased similarity of neural responses derived from the Sharpening model. Changes in functional and effective connectivity provided the most direct support for the Synchrony model, with increased coupling from right temporoparietal cortex to the anterior cingulate cortex for OLD relative to NEW objects, particularly for the strong primeable condition. This occurred despite no change in the activity levels in right temporoparietal cortex and decreased activity for OLD objects in the anterior cingulate cortex, demonstrating increased impact of activity of one region on the other that is consistent with enhanced synchronization of activity. Furthermore, these changes were correlated with inter-participant variability in behavioral priming. Top-down negative coupling that increased with repetition, a central prediction of the Predictive Coding model, was not observed among any regions, with only positive coupling observed, and changes in coupling were found to be largely independent of repetition suppression magnitudes. With respect to the Facilitation model, neural responses were indeed found to peak at earlier times for OLD compared to NEW objects in regions showing repetition suppression, although these effects failed to interact with primeability. While peak response amplitude, as opposed to timing, was found to interact with primeability in the left fusiform gyrus, it failed to correlate with priming magnitudes across participants. With respect to the Sharpening model, the spatial similarity of neural responses assessed with MVPA interacted with primeability in a manner qualitatively similar to behavioral priming effect sizes. However, these effects were strongly related to the average response amplitude and estimates of the signal-to-noise ratio in each region, largely eliminating differences in similarity after adjustment for these factors.

On the whole, the results provide the strongest support for the Synchrony model. None of the other models exhibited robust correlation with both within- and between-participant measures of behavioral priming magnitude. However, even this model failed to provide an explanation of repetition suppression magnitude. Despite the typical joint observation of repetition suppression and repetition priming, studies have often failed to observe robust intercorrelation of these two phenomena across participants and/or trials (e.g. McMahon & Olson, 2007; Race et al., 2009; Salimpoor et al., 2010; Sayres & Grill-Spector, 2006; Xu et al., 2007), with the most frequent reports of correlations in lateral prefrontal cortex (e.g. Bunzeck et al., 2006; Dobbins et al., 2004; Horner & Henson, 2008; Maccotta & Buckner, 2004; discussed in Horner, 2012). In the current study, the phenomena were largely independent of each other across participants, despite extremely robust effects for both phenomena (see Figures 3, 4, and 7) and more than 3 times the number of participants in most task-based fMRI studies (often 20 or fewer). Consistent with this, correlations between effective connectivity changes and priming magnitudes were of equivalent size after partialling repetition suppression magnitudes. Our results suggest that repetition suppression and repetition priming reflect the operation of at least partially separate mechanisms, consistent with prior studies that have experimentally decoupled the effects either with TMS (e.g. Wig et al., 2005) or with changes in task between study and test phases (e.g. Dobbins et al., 2004; Race et al., 2009; Horner & Henson, 2008; 2012). The functional role of repetition suppression thus remains unclear. It is possible that repetition suppression instead plays a more direct role in optimizing energy use and efficiency, without making specific functional contributions to cognition.

The failure to observe robust support for the Predictive Coding, Facilitation, and Sharpening models may be at least partially due to methodological factors. The current study was conducted with fMRI, which afforded simultaneous testing of all four major theoretical models. However, fMRI utilizes the BOLD signal rather than the underlying electrophysiological signals, and BOLD fMRI may be blind to effects that occur at the more rapid time scale of milliseconds to hundreds of milliseconds. For example, in a recent study of short-term priming in picture naming using electrocorticography (or ECoG), we found evidence of both earlier neural response onset consistent with the Facilitation model, as well as increased top-down effective connectivity from left frontal cortex to left ventral temporal cortex with repetition within the first 200 milliseconds of processing (Korzeniewska et al., 2020), potentially consistent with the Predictive Coding model. While this study examined only short-term effects (less than 20 seconds separating repetitions) and did not have the statistical power to evaluate the relationship between changes in coupling, repetition suppression magnitude, and behavioral priming magnitude across participants, the observation of effects over a much more rapid physiological time scale suggests the need for further studies with higher temporal resolution (such as ECoG, MEG, and joint fMRI/EEG studies). It is similarly unclear whether spatial correlations in fMRI are a sufficient test of the Sharpening model (e.g. Alink et al., 2018; Ramirez, 2018). However, multiple other studies have tested the Sharpening model using a range of methods in both monkeys and humans, with little or no supportive evidence to date at the most common time scales used in priming studies (i.e. within a daily session; Alink et al., 2018; De Baene & Vogels, 2010; Gotts et al., 2015; Kaliukhovich et al., 2013; Li et al., 1993; McMahon & Olson, 2007; Miller et al., 1993; Weiner et al., 2010; see Gotts, 2016, for discussion).

The detection of increased functional coupling between right temporoparietal cortex and the anterior cingulate with repetition is novel to the current study. Despite its robustness across separate participants performing covert versus overt naming (see Figure S2), as well as its intercorrelation with priming magnitude, it is still unanticipated by previous studies. Rather than viewing this connection in isolation, it is useful to examine its context within the effective connectivity model that included interactions with portions of the striatum (right putamen), STG and the right fusiform gyrus (Figure 6A). It is possible that this entire circuit functions to retrieve prior stimulus-responses associations encoded through cortico-striatal loops (Atallah et al., 2007; Boettiger & D’Esposito, 2005; Packard & Knowlton, 2002; Seger & Miller, 2010; see also Henson et al., 2014; 2017; Race et al., 2019; Suzuki, 2008). Some evidence consistent with the Synchrony model that we previously observed in MEG in covert picture naming was also right lateralized, with enhanced evoked power in the theta/alpha frequency ranges involving the right fusiform gyrus and right lateral prefrontal cortex (Gilbert et al., 2011). Other studies have highlighted a role for the right temporoparietal cortex in both attention- and memory-related contexts as a portion of the ventral attention network (e.g. Cabeza et al., 2011; 2012; Corbetta & Shulman, 2011). Nevertheless, it will be important for future studies, both physiological and neuropsychological, to further examine the role of these regions in repetition priming and other memory-related contexts.

On a final note, the lack of interaction with task (overt versus covert naming) for the effects in the current study suggest that studies with limited overt verbal responding may be used productively in fMRI, provided that there is a way to account for the increased artifacts that are expected to accompany this form of responding. Here, we included a non-speech control and covaried the global level of correlation present in each condition for each participant, and results surviving those controls were found to correlate with behavioral performance (see also Gilmore et al., 2019; Jasmin et al., 2019). Similarly, the employment of a slow event-related rather than a more typical rapid event-related fMRI design allowed us to separate the temporal contour of evoked responses from estimates of functional and effective connectivity, as well as permitting the interleaving of different trial types and restricting connectivity estimates to correct trials. These tools should be applicable to exploring task-based connectivity effects in a variety of cognitive domains, complimenting approaches already in use in resting-state fMRI research.

## Conclusion

We set out to test the major models posited to explain the neural bases of repetition priming. Our findings were most supportive of the Synchrony model. The strength of coupling between specific brain regions was correlated with the strength of priming. However, these coupling changes were unrelated to repetition suppression effects. Indeed, none of the models we tested provided insight into the relationship between repetition priming and repetition suppression, thereby suggesting that these effects may depend on at least partially distinct mechanisms.

## Acknowledgements

We thank Ally Ossowski for assisting with data collection and analysis, Gang Chen and Daniel Glen for assisting with the structural equation modeling analyses, Vinai Roopchansingh for aid in designing the sagittal fMRI scanning sequences, and Adrian Gilmore, Michal Ramot, Andrew Persichetti, Jason Avery, Al Braun, Bob Cox, Peter Bandettini, Fernando Ramirez, Eli Merriam, and Chris Baker for helpful discussions. This study was supported by the National Institute of Mental Health, NIH, Division of Intramural Research (ZIAMH002920; ClinicalTrials.gov ID NCT00001360). The funders had no role in the study design, data collection and analysis, decision to publish, or preparation of the manuscript.

## Author Contributions

Conceived and designed the experiments: SJG, AM

Performed the experiments: SJG, SCM

Analyzed the data: SJG, SCM

Wrote the paper: SJG, AM

## STAR Methods

### Lead Contact and Materials Availability

#### Lead Contact

Further information and requests for resources and reagents should be directed to and will be fulfilled by the Lead Contact, SJG.

#### Materials Availability Statement

This study did not generate new unique reagents.

### Experimental Model and Subject Details

#### Ethics Statement

Ethics approval for this study was granted by the NIH Institutional Review Board (protocol 93-M-0170, clinical trials number NCT00001360).

#### Participants

Thirty-two participants performed the Overt Naming Task (18 females) with a mean (SD) age of 24.03 (3.58) years (range: 19 to 38), and 28 additional participants performed the Covert Naming Task (19 females) with a mean (SD) age of 23.43 (1.62) years (range: 21 to 28). Participants were right-handed, neurologically healthy native English speakers with normal or corrected-to-normal vision. All participants granted informed consent and were monetarily compensated for their participation.

### Method Details

#### Experimental Stimuli

Participants completed either overt or covert picture naming, consisting of 200 colored photographic images of animals, plants, foods, and everyday objects. Images were presented across two lists of 100 pictures each, with the lists matched in conceptual category membership and in average lexical properties of picture names (omnibus *F* statistics all < 1), including lexical decision times on the names (mean RT = 632.3 ms, SD = 71.9 ms) and log HAL frequency determined by the English Lexicon Project database (mean = 8.57, SD = 1.54) (Balota et al., 2007). Images were resized to 600 × 600 pixels and presented against a gray background (RGB value: 75, 75, 75). Outside of the scanner, pictures presented on a laptop subtended approximately the central 6° x 5° of visual angle (horizontal X vertical). Inside of the scanner, pictures subtended approximately the central 7.8° X 6.2° of visual angle (horizontal x vertical).

#### Naming Tasks

Participants in both Overt and Covert Naming conditions initially named one set of 100 images aloud three times through in a pseudorandom order outside the scanner (in a quiet testing room). In each naming trial, the trial started with a central fixation cross for 500 ms, followed by the picture to be named for 200 ms. The picture offset was followed by a blank screen for 1300 ms, yielding a total trial duration of 2000 ms. Participants were instructed to name each image aloud as quickly and accurately as possible, with correct performance and error responses notated by the experimenter, and response time marked according to voice onset using a microphone on the display computer (Presentation software package Version 11.3, www.neurobs.com).

After a delay of approximately 30 minutes, participants performed either Overt or Covert Naming inside the MR scanner. In both tasks, a naming trial consisted of a central black fixation cross presented for 1000 ms, followed by the picture to be named for 300 ms, followed by a blank screen for a period ranging from 6700 to 12700 ms at multiples of the TR (6700, 8700, 10700, 12700 ms) and sampled with a uniform distribution (Figure 2; see Bandettini & Cox, 2000, for discussion of optimal ISIs in slow event-related fMRI designs). For Overt Naming, participants spoke into an MR-compatible microphone placed next to the head coil approximately 3-5 cm from the participant’s mouth. For Covert Naming, participants were instructed to name the pictures silently to themselves, pressing a response button to mark the beginning of their naming response. In both tasks, the 100 pictures named pre-fMRI (OLD) were randomly intermixed with 100 pictures that were novel for the fMRI session (NEW), with trials organized into 5 runs of 40 pictures each. For Covert Naming, participants completed an additional overt naming session of all 200 pictures immediately after fMRI, in order to confirm that their button press response times during fMRI agreed with actual overt response times (same timing and methods used as for the pre-fMRI session). Experimental lists (OLD versus NEW) were counterbalanced across participants, and only correct trials were included in priming estimates and task analyses.

#### Recording Naming Responses during MRI

Spoken responses were captured with an Opto-Acoustics FOMRI-III NC MR-compatible microphone with built-in noise cancellation and routed into an M-Audio FastTrack Ultra 8-R USB audio interface. Responses were recorded with Adobe Audition. To calculate response times, the stimulus presentation computer emitted a square wave pulse at the onset of each trial and a custom Matlab program calculated the time difference between the square pulse onset and voice response onset for each trial.

#### MRI Methods

Images were acquired with a General Electric Signa HDxt 3.0T scanner (GE Healthcare) using an 8-channel receive-only head coil. A high-resolution T1-weighted anatomical image (MPRAGE, magnetization-prepared rapid gradient-echo) was obtained for each participant (124 axial slices, 1.2 mm slice thickness, field of view = 24 cm, 224 × 224 acquisition matrix). Functional (T2*-weighted) images were acquired using a gradient-echo echo-planar imaging (EPI) sequence [Array Spatial Sensitivity Encoding Technique, ASSET, acceleration factor = 2, TR = 2000 ms, TE = 27 ms, flip angle = 60°, 40 sagittal slices (3.5 mm slice thickness), field of view = 216 mm, 72 × 72 acquisition matrix, voxel resolution = 3.5 × 3.0 × 3.0 mm^3^]. Each experimental task run lasted 7 min 40 sec for a total of 230 consecutive whole-brain volumes, with each participant receiving a total of 5 runs. Foam earplugs were worn by participants to attenuate scanner noise and participants’ head positions were stabilized using foam pillows. All EPI data were evaluated for transient head motion artifacts, with included scans required to be less than or equal to 0.3mm/TR using AFNI’s @1dDiffMag function (comparable to mean Framewise Displacement, Power et al., 2012). Independent measures of cardiac and respiration cycles were recorded during the task scans for later removal.

#### fMRI Data Preprocessing

Preprocessing utilized the AFNI software package (Cox, 1996), applying steps in the following order: 1) removal of the first 3 TRs to allow for T1 equilibration; 2) 3dDespike to bound outlying time points per voxel within 4 standard deviations of the time series mean; 3) 3dTshift to adjust for slice acquisition time within each volume (to t=0); 4) 3dvolreg to align each volume of a run’s scan series to the first retained volume of the first run; 5) each scan was then spatially blurred by a 6-mm Gaussian kernel (full width at half maximum) and divided by the voxelwise time series mean to yield units of percentage signal change. De-noising of each scan then utilized the ANATICOR nuisance regression approach (Jo et al., 2010; see also Gotts, Simmons, Milbury, et al., 2012). White matter and large ventricle masks were created from the aligned MPRAGE scan using Freesurfer (e.g. Fischl et al., 2002), and a large draining vein mask was created from a standard deviation map of the volume-registered EPI data (from step 4 above). All masks were resampled to EPI resolution and eroded by 1 voxel to prevent partial volume effects with gray matter voxels, and the related nuisance time series were calculated on the volume-registered data just prior to spatial blurring (after step 4 and prior to step 5 above). Nuisance regression for each voxel was performed on the spatially blurred volume-registered data (after step 5 above), and the regressors consisted of: 6 head-position parameter time series (3 translation, 3 rotation), 1 average eroded ventricle time series, 1 “localized” eroded white matter time series (averaging the time series of all white matter voxels within a 20-mm radius sphere), 1 eroded draining vein time series, 8 Retroicor time series (4 cardiac, 4 respiration) calculated from the cardiac and respiratory measures taken during the scan (Glover et al., 2000), 5 Respiration Volume per Time (RVT) time series to minimize end-tidal CO_2_ effects following deep breaths (Birn et al., 2008), and the first 3 principal component time series calculated on a union mask of the nuisance tissues (white matter, ventricles, draining veins) (aCompCor regressors: Behzadi et al., 2007; Stoddard et al., 2016). Prior to regression, all nuisance time series were detrended by a 4^th^-order polynomial function to remove slower scanner drift and drift in head position, with the de-noised residuals detrended in the same manner during regression. After regression, de-noised residual time series were transformed to standardized anatomical space (Talairach-Tournoux) for task analyses at a resolution of 3 mm^3^ isotropic.

### Quantification and Statistical Analysis

#### Normative Analyses of Naming Response Times and Primeability

A within-participant estimation of “primeability” (Strong versus Weak) in object naming was determined from normative analyses of response times across all 60 participants. Response times on correct naming trials were averaged across participants when encountered for the first time as NEW objects (i.e. the first presentation in the pre-fMRI session, first presentation during the fMRI session, or first presentation in overt naming for the post-fMRI session for Covert Naming participants), as well as for OLD objects (i.e. presentation during fMRI of objects seen during the pre-fMRI session; for Covert Naming participants, post-fMRI response times to OLD objects were used). Only response times in overt naming sessions were used for this purpose, since button press response times during Covert Naming were systematically faster than for overt naming sessions by as much as 200-300 ms per participant. Test-retest reliability of mean NEW response times per object was assessed across Overt and Covert Naming experiments using Pearson correlation, as well as the test-retest reliability of priming magnitude per object, estimated as the difference in response time when an object was OLD versus NEW. Reliability of NEW responses and priming magnitudes across Overt and Covert Naming experiments was high for both measures, but was highest for NEW responses [r(198)=.704 vs r(198)=.436]. Given the high reliability of mean NEW response times across experiments, as well as the strong relationship between mean NEW responses and priming magnitude by item when combining experiments (larger priming for slower NEW responses; Figure 4B), “primeability” was calculated as a median split of mean NEW response times, with Strong Primeable objects defined as the 100 objects with the slowest mean NEW response times across participants and Weak Primeable objects defined as the 100 objects with the fastest mean NEW response times across participants.

#### Behavioral Priming Analyses

Response times to correct naming trials during fMRI were tabulated for each participant, separately for each experimental condition [Repetition (OLD, NEW) crossed with Primeability (Strong, Weak) for a total of 4 conditions]. Repetition priming magnitudes per participant were then calculated for Strong and Weak Primeable conditions using effect size (Cohen’s *d*), the difference in means (NEW – OLD) divided by the pooled standard deviation. For purposes of correlating priming magnitudes with measures of neural activity derived from fMRI, effect sizes determined for Covert Naming participants were averaged from the fMRI and post-fMRI sessions (during which the same items were named overtly).

#### fMRI Task Analyses

Traditional task analyses were conducted at the voxel level using a general linear model (GLM), in which the data at each time point are treated as the sum of all effects thought to be present at that time point, plus an error term. Responses associated with each condition were modeled using TENT basis functions in AFNI, with a separate regressor for each time point following the stimulus onset, permitting empirical estimation of the hemodynamic response function (HRF) shape. This approach assumes that all responses for a given condition share the same response shape, but makes no assumption as to what the shape of that response might be. Responses for each participant were modeled over 8 time points from *t*=0 seconds to *t*=14 seconds in increments of the TR (2 seconds). One additional regressor of no-interest was coded for error trials in naming (either omissions or commissions). For the purposes of statistical testing, peak response magnitudes were estimated by averaging the 3^rd^ and 4^th^ time points of the TENT function corresponding to the peak of the typical BOLD response, reflecting activity 4 to 8 seconds post-stimulus onset; visual examination of the beta coefficients confirmed that this period did indeed represent the typical peak well (see Figure 7). Single-participant contrasts for overall stimulus response (pooling conditions) and effect of repetition (either repetition suppression or enhancement, pooling across Strong/Weak Primeability conditions) were tested using linear tests within the GLM. Beta coefficients from the regression for each participant were submitted to group-level analyses of the overall stimulus response and effects of repetition.

A group-level effect of Stimulus Response relative to baseline was evaluated in each voxel separately for Overt and Covert Naming using a one-sample t-test across participants of the related beta coefficients (β) versus 0 (averaging the 3^rd^ and 4^th^ TR β’s at the peak response), with positive β’s indicating above-baseline and negative β’s indicating below-baseline responses. Two statistical thresholds were examined: 1) a minimum level of significance at P<.05 when pooling participants across the two tasks, corrected for whole-brain comparisons using false discovery rate (FDR) to *q*<.05 (e.g. Genovese et al., 2002), and 2) a stringent level of significance in each task individually (P<.0001, *q*<.00016); given the number of voxels meeting the threshold in the brain volume (6322 voxels at q=.00016 in Covert Naming, 9287 voxel at q=.00009 in Overt Naming), this level of FDR corresponds roughly to the expectation of only 1 false positive voxel in each task. At the more stringent threshold (with the extremely low FDR), responses can be said to replicate across tasks.

A similar approach was taken in evaluating Repetition effects (OLD versus NEW objects) at the group level, collapsing across Strong/Weak Primeability conditions. Two statistical thresholds were examined using a paired t-test (OLD-NEW) across participants: 1) a minimum level of significance (P<.05, q<.05) when pooling participants across the two tasks, and 2) a stringent level of significance in each task individually (P<.00001, q<.00006), with an expectation of less than 1 false positive voxel in each task.

#### fMRI Task-Based Functional Connectivity Analyses

In order to avoid contamination of task-based functional connectivity estimates from the temporal contour of the evoked responses of individual trials, only the average peak BOLD response was retained from each individual trial (the raw average of the 3rd and 4th timepoints of the task residual timeseries following the onset of each stimulus). For these analyses, data had undergone preprocessing with nuisance regression (described above), as well as having removed the condition-level mean responses during GLM regression analyses. This resulted in an “item series” of peak BOLD responses from correct trials in each individual voxel, with a maximum length of 100 OLD and 100 NEW trials and each divided approximately in half for Strong and Weak Primeability conditions (e.g. a maximum series of approximately 50 Strong Primeable, OLD trials). Functional connectivity analyses utilized a 2×2×2 mixed effects design with Task (Overt, Covert) as a between-participant variable, Repetition (OLD, NEW) and Primeability (Strong, Weak) as within-participant variables and participant as a random variable. For all analyses, two sets of tests were of primary interest: 1) a main effect of Repetition (OLD versus NEW) collapsing across Primeability, and 2) an interaction of Repetition X Primeability, given that this is the pattern observed in behavioral priming with larger differences between NEW and OLD response times for Strong Primeable compared to Weak Primable conditions (Figure 4). Accordingly, the familywise alpha for multiple-comparisons correction was set at P<.025 (.05/2) in order to correct for two sets of voxelwise (or ROI-level) tests. Our functional connectivity analyses employed the 3 steps used previously by Gotts and colleagues for whole-brain functional connectivity analyses: seed definition, target ROI selection, and region-to-region correlation analysis (e,g., Berman et al., 2016; Gotts, Simmons, Milbury, et al., 2012; Jasmin et al., 2019). We undertook all 3 steps for both the Repetition main effect and the Repetition X Primeability interaction effect. The further interaction of these primary effects with Task was also evaluated to examine the possibility that any results could simply be due to the presence of speech artifacts (e.g. present in Overt but not Covert Naming).

Seeds were identified using whole-brain “connectedness” (e.g. Cole et al., 2010; Gotts, Simmons, Milbury, et al., 2012; Salomon et al., 2011). The average Pearson correlation of each voxel’s “item series” (the array of peak BOLD responses to individual trials of the same type) with the item series in all voxels responding above baseline in the task (P<.0001, q<.05, for each individual participant) was calculated to create a 3D reduction of the 4D (3D + Time) dataset for each condition. In other words, the correlation of a particular voxel’s responses was calculated with those of all task-responsive voxels, storing the average of those correlations back into the voxel. Since connectedness in this case reflects the average level of correlation with task responsive voxels, it gives an indication of how intercorrelated a given voxel is with those voxels most engaged by the task. This approach, akin to centrality in graph theory, has been used previously in studies of both resting-state (e.g., Berman et al., 2016; Cole et al., 2010; Gotts, Simmons, Milbury, et al., 2012; Meoded et al., 2015; Salomon et al., 2011; Smith et al., 2019; Stoddard et al., 2016; Watson et al., 2019) and task-based functional connectivity (e.g., Jasmin et al., 2019; Song et al., 2015; Steel et al., 2016). Linear mixed effects (LME) models (using AFNI’s 3dLME, Chen et al., 2013) were constructed whose dependent variables were the voxel-wise connectedness maps in each experimental condition. Task, Repetition, Primeability, and their interactions were included as fixed effects. The global level of correlation among all brain voxels, GCOR (Saad et al., 2013), was included as a nuisance covariate in order to model any residual motion and/or breathing artifacts present after the nuisance regression (see Gotts et al., 2020, for discussion). Participant was treated as a random intercept. Cluster-size correction was used to control the Type I error rate. The average smoothness of the de-noised functional time series was estimated with AFNI’s 3dFWHMx, using the empirical, spatial autocorrelation function (June 2016). 3dClustSim (June 2016) was then used to run a Monte Carlo simulation with 10,000 iterations in a whole-brain mask in Talairach space within which the analyses were performed. Importantly, the smoothness estimates and noise simulations did not assume Gaussian distributions of activity, which has been shown to inflate the false positive rate in studies using more traditional cluster size correction (e.g. Cox et al., 2017; Eklund et al., 2016). Clusters were selected at a cluster defining threshold of P<.001, familywise alpha of P<.025 (.05/2, for two sets of tests), minimum cluster size k=25 voxels.

The seed definition step was then followed with more typical seed-based correlation analyses. The item series within each seed region was averaged across voxels to form ROI-averaged item series, which were correlated with the item series for every voxel in the brain, separately for each experimental condition. These correlations were Fisher z-transformed and used as dependent variables in LME models with the same fixed and random effects as for the seed detection step. We tested for the main effect of Repetition and the Repetition X Primeability interaction at a voxel threshold of p<.001, with correction by cluster size for whole-brain comparisons as well as the number of seeds tested (i.e. FWE correction to P<;[.025 / (number of seeds)]. Results were then further masked by voxels exhibiting above-baseline responses in both tasks individually (P<.05, q<.05), leaving clusters of voxels that both showed changes in functional connectivity with stimulus repetition and that were engaged in the task. Secondary target regions were then combined together with seed regions, as well as any regions exhibiting repetition suppression in both tasks, to arrive at a full set of ROIs.

Regions were sampled as 6-mm-radius spheres centered on the peak statistic used to identify each ROI (F-statistic from the LME on connectedness or seed-based correlation tests for ROIs showing changes in functional connectivity; the t-value of the OLD versus NEW comparison of beta coefficients for repetition suppression regions). Region-by-region matrix analyses were then conducted using the same LME approach applied to connectedness and the seed-based tests, allowing the examination of all inter-regional relationships. Multiple comparisons in the region-by-region analyses were controlled with FDR (q<.05).

#### fMRI Task-Based Effective Connectivity Analyses

Single-trial responses (same data as for functional connectivity analyses) from the 7 regions identified in the whole-brain functional connectivity analyses (including repetition suppression regions) were submitted to effective connectivity analyses (e.g. Friston, 1994; 2011; Smith et al., 2012). Of the various effective connectivity approaches (e.g. Dynamic Causal Modeling - DCM, Granger Causality, Multivariate Autoregressive Modeling, etc.), Structural Equation Modeling, or SEM (e.g. Chen et al., 2011; McIntosh & Gonzalez-Lima, 1994; Price et al., 2009), provided the best match for the current data characteristics. In particular, any approaches requiring block designs (e.g. DCM) and/or the use of time series (e.g. Granger-based methods) were not appropriate, since the data consisted of a series of peak BOLD responses that had been notched out of the original time series in an event-related design with interleaved trial types. As with other effective connectivity approaches, we first performed a search for the model that best accounted for the pattern of covariance among the ROIs when pooling data across all conditions and participants (using AFNI’s 1dSEM, Chen et al., 2011). A search was performed with both tree growth and forest growth algorithms using the Akaike Information Criterion (AIC; Akaike, 1973), a measure of out-of-sample prediction error, to choose among different SEM models. The optimal model had 11 directional connections among the ROIs.

Following the model search step, the optimal model was parameterized for each participant’s condition-specific datasets individually (AFNI’s 1dSEMr, Chen et al., 2011). The parameterization failed to converge for two datasets (the NEW, Weak Primeable condition for one participant, the OLD, Strong Primeable condition for another; both Covert Naming participants), and these sets were excluded from further analyses. The results of the successful parameterizations were then submitted to an LME analysis that paralleled the one performed on the functional connectivity data, with the SEM parameters as the dependent variable and Task, Repetition, Primeability, and their interactions included as fixed effects. GCOR was included as a nuisance covariate, and Participant was treated as a random intercept. One out of the 11 SEM parameters (from the R TP to the Left Frontal ROIs) failed to differ from zero across participants overall or in any individual condition and was excluded from further analyses. The remaining 10 parameters were tested for overall effects of Repetition and Repetition X Primeability interactions, as with the functional connectivity analyses, with multiple comparisons corrected by FDR (q<.05). Further interactions with Task (Overt, Covert) were also evaluated. For connections exhibiting a significant Repetition X Primeability interaction, follow-up contrasts were conducted to clarify the nature of the interaction, comparing parameters to OLD and NEW objects separately for Strong versus Weak Primeable conditions, with multiple comparisons corrected by FDR (q<.05).

#### Evaluating Timing Predictions of the Facilitation Model

A central prediction of the Facilitation Model is that neural responses should be temporally advanced for OLD compared to NEW objects in regions showing repetition suppression, with an earlier peak response (e.g. James et al., 2000; James & Gauthier, 2006). In order to examine this prediction for the evoked responses in fMRI, we fit a simple gamma variate function to the TR-specific beta regression coefficients calculated during the GLM analyses in the four repetition suppression ROIs, with a separate response function for each of the four experimental conditions [Repetition (OLD,NEW) X Primeability (Strong,Weak)]:

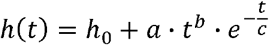

*h*(*t*) represented the hemodynamic response as a function of time *t*, with 4 modifiable parameters: *h*_0_, representing a baseline value of *h*; *a*, a scaling parameter on the height of the curve; *b*, an exponent on *t* determining the rise time of the curve; and *c*, a time constant for the exponential decay of the curve. The time of the peak response (*t*_*peak*_) then corresponded to the value of *t* at which the derivative of *h*(*t*) with respect to t equaled zero:

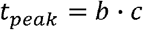

The peak amplitude, or height of the response, corresponded to the value of *h* at *t*_*peak*_ For each curve, the best-fitting parameter values for all 4 free parameters were determined initially by a course grid search, followed by fine-tuning through a gradient descent error minimization until stability had been reached (<;=300 iterations). The estimates of *t*_*peak*_ and the peak amplitude were then submitted to separate LME analyses as the dependent variables, including Task, Repetition, Primeability, and their interactions as fixed effects and Participant treated as a random intercept. Multiple comparisons were corrected by FDR (q<.05).

#### Evaluating Predictions of the Sharpening Model with MVPA

The Sharpening Model predicts that OLD objects should have spatial patterns across repetition suppression ROIs that are less similar to one another than for NEW objects, consistent with the loss of neural overlap among the neural representations. To evaluate this, we applied multi-voxel pattern analysis (MVPA) to the single-trial BOLD responses in each experimental condition (OLD, NEW X Strong, Weak Primeable). For these analyses, the data used were post-nuisance regression in preprocessing but prior to the removal of condition means in GLM analyses. Each single-trial response was calculated as the average of the 3^rd^ and 4^th^ TRs post-stimulus onset (the 4-6 sec and 6-8 sec TRs), i.e. the peak response, minus the average of the 1^st^ TR in the trial (0-2 sec) and the TR previous to it (−2-0 sec), i.e. the baseline response. This resulted in a voxelwise pattern of BOLD responses within an ROI. For a given participant and ROI, the spatial correlation (Pearson) of all trials with all trials within the same condition (e.g. NEW, Strong Primeable) was calculated across voxels, storing the median correlation value to protect against skewing effects. The median correlations then served as the dependent variable in an LME analysis, including Task, Repetition, Primeability, and their interactions as fixed effects and Participant treated as a random intercept. Multiple comparisons were corrected by FDR (q<.05). Further interactions with Task (Overt, Covert) were also evaluated. For ROIs exhibiting a significant Repetition X Primeability interaction, follow-up contrasts were conducted to clarify the nature of the interaction, comparing median correlations to OLD and NEW objects separately for Strong versus Weak Primeable conditions, with multiple comparisons corrected by FDR (q<.05).

The use of individual trial-level and voxel-level BOLD responses was expected to suffer from poor signal-to-noise ratios when scanning at 3 Tesla, with a high potential for “attenuation” of spatial correlation values due to noise (e.g. Spearman, 1904). This could be particularly problematic when the ROIs of interest have known differences in overall activity levels in different conditions, in this case due to repetition suppression – which is how the ROIs were defined. If OLD objects have BOLD responses that are closer to the baseline level than NEW objects, the spatial correlation values could be closer to 0 simply due to lower beta coefficients across the ROIs and poorer signal-to-noise ratios. In order to examine this issue, two nuisance covariates were included in a subsequent LME analysis of each ROI: 1) for average beta coefficient across the ROI, and 2) for estimated signal-to-noise ratio of trials in each condition for each participant. The signal-to-noise ratio (SNR), defined as the variance of the signal divided by the variance of the noise, was estimated for a given condition in the following manner: 1) for each trial, the difference between the peak and baseline TRs represented the single-trial BOLD response; the variance of this pattern across voxels served as the numerator in the SNR calculation, 2) a noise estimate for each trial was calculated by randomly selecting two baseline patterns from the same ROI and condition, subtracting one from the other, and then calculating the variance of this pattern across voxels, 3) dividing the variance in step 1 by the variance in step 2 yielded the SNR estimate for that trial, 4) the median SNR value across trials in each condition served as the second nuisance covariate in the new LME analysis. It is important to mention that the estimate of the peak responses in each trial is not noiseless in this case, so this estimate of SNR is expected to be an upper bound on the true SNR. A strong fit of the two nuisance covariates to the ROI data, combined with a change in significance levels from the LME analysis without the covariates, would indicate results consistent with correlation attenuation. These relationships were further evaluated by examining scatterplots of the nuisance variables on the x-axis and median spatial correlation level on the y-axis, overlapping results across the different experimental conditions.

### Data and Code Availability

Data are available via the XNAT platform. Users will need to request access through the XNAT system. This can be done by creating an XNAT user account and pressing the ‘request access’ link. All analysis code used in the paper is available on request.

## Supplemental Information Titles and Legends

**Table S1. Regions of Interest.** Regions of interest used in Functional (FC) and Effective Connectivity (EC) Analyses (Repetition Suppression, RS, and FC regions), as well as subsequent tests of Facilitation and Sharpening models (RS regions only). Standard space coordinates (Talairach-Tournoux) of the peak statistic used to identify effects are reported, along with the spatial extent of the associated clusters by effect type. For Right Fusiform and ACC ROIs, detected in both RS and FC analyses, the peak coordinate used is related to the peak of the FC effects within the voxels that overlapped across the two effects (in order to maximize the observation of any FC/EC effects). Full ROI-ROI matrix analyses of FC/EC effects, as well as tests of the Facilitation model, used spherical ROIs centered on the peak coordinates. MVPA tests of the Sharpening model used the full spatial extent of RS clusters. Related to Figure 5.

**Figure S1. Individual Picture Naming Response Times are Highly Variable but Become Reliable when Averaged.** Test-retest reliability of individual response times (RT) was evaluated using the pre-fMRI Overt Naming sessions for participants performing both Overt (N=32) and Covert Naming (N=28) conditions during fMRI. (A) Each participant named a set of 100 pictures presented 3 times in a pseudorandom order in a quiet testing room prior to fMRI (Repetitions 1-3). As expected, decreased RTs were observed on the 2^nd^ and 3^rd^ repetitions relative to the 1^st^ presentation in both sets of participants (calculated on correct naming trials). Error bars indicate standard error of the mean (SE). (B) Test-retest reliability (Pearson *r*) was calculated for each participant using the 3 pre-fMRI repetitions (1^st^ vs 2^nd^, 2^nd^ vs 3^rd^, 1^st^ vs 3^rd^), including only items named correctly for all 3 repetitions (a mean of 80.48 items per participant out of a possible 100). The distributions of mean reliability (over all combinations of repetition pairs) across all participants is shown for Overt and Covert Naming participants separately using boxplots, along with all individual participant datapoints as open circles. The red horizontal line in each box plot represents the median (50th %-ile), the blue horizontal lines just above and below the median represent the 25th and 75th %-iles, the black horizontal lines enclose +/− 2.7 standard deviations of the mean (99.3% coverage of a normal curve), and the boundaries of the horizontal notches inside the 25th and 75th %-iles depict the 95% confidence limits of the median. The distributions do not differ significantly between the Overt and Covert Naming participants (median Covert reliability = 0.1607; median Overt reliability = 0.1350; overall median = 0.1567). (C) Despite the poor test-retest reliability of the individual RTs, averaging RTs across participants by item (or across items within-participant; see Figure 4C) leads to large improvements in reliability. Shown is the test-retest reliability (Pearson *r*) when averaging across participants in each task condition by item (Overt vs Covert). As pre-exposed sets were counter-balanced across participants, averaging included slightly less than half of the number of participants in each condition (correct responses only; median number of participants averaged per item was 15 for Overt Naming, 11 for Covert Naming). After averaging over participants and only utilizing RTs from Repetition 1 (the first time each participant encountered an item), the test-retest reliability of item RTs increases to 0.704. These results suggest that successfully identifying the neural correlate of repetition priming will require a degree of trial averaging/pooling; i.e. measuring the change in RT on individual trials and relating that to the change in single-trial BOLD responses will not be practical, even if only considering the reliability of behavior. Related to Figure 4.

**Figure S2. Agreement of Repetition X Primeability Interaction in Seed Detection Across Tasks.** The Repetition X Primeability interaction in whole-brain connectedness found in the right temporoparietal (R TP) cortex failed to interact significantly with Task (Overt vs Covert Naming). Shown in the left panels are voxels that are significant in each task individually (P<.05, uncorrected) (blue = Overt Naming only; yellow = Covert Naming only; red = significant in both). These results were masked by voxels that showed a consistent interaction pattern across both tasks (P<.05, uncorrected, when including data from both tasks; minimum cluster size of 100 voxels). Virtually all of the voxels in the R TP ROI surviving whole-brain correction in the full data (Figure 5A, shown for reference in right panels) were significant at an uncorrected level in both tasks individually. A cluster in the right frontal cortex corresponded to a cluster in the full analysis that failed to survive whole-brain correction, but showed a similar pattern to that in R TP. The locations of both the R TP and right frontal clusters agree well with the locations of clusters in the Ventral Attention network (e.g. Yeo et al., 2011). Related to Figure 5.

**Figure S3. Increased coupling occurs with no increase in activity levels.** The increased coupling observed from the R TP to ACC ROIs, particularly in the Strong Primeable condition (see bottom panel), happens despite no increase in activity levels in R TP (Strong Primeable, OLD vs NEW: *P*>.77) and a decrease in activity in ACC (Strong Primeable, OLD < NEW: *P*<1.5×10^−13^). In the absence of a change in the synchronization of the underlying activity in R TP and ACC, increased synaptic coupling onto ACC would be expected to lead to higher, not lower activity in ACC. The fact that activity is not increased in R TP, coupling from R TP is increased to ACC, and activity is decreased in the ACC is most consistent with increased synchronization of neural activity one or both regions simultaneously with the increased interregional coupling (see Gotts, 2003, for related computational modeling and discussion). The repetition suppression in the ACC is itself largely unrelated to the coupling increases, as detailed in the partial correlation analysis in the main text. Related to Figure 6.

**Figure S4. Similarity measures calculated on single-trial BOLD responses are strongly related to average activity level and signal-to-noise ratio.** Evaluation of the Sharpening model using MVPA involves calculating spatial similarity measures across voxels and within conditions that differ in average activity level (identified as showing RS in the beta coefficients). In the current experiment, individual pictures were presented only once during fMRI, requiring the use of single-trial responses in MVPA, which are expected to be noisy. While the raw measures of similarity follow the predictions of the Sharpening model, particularly for the L Frontal and ACC ROIs (Figure 8A), they also follow the pattern of the average beta coefficients (averaged across voxels within an ROI). After adjustment for average beta coefficients and estimated signal-to-noise ratio (SNR), virtually all of the MVPA effects become non-significant (Figure 8B). (A) The dependence of similarity (Pearson *r*; on y axes) on beta values and estimated SNR (x axes) are shown more directly using scatterplots. In each scatterplot, each participant contributes 4 datapoints, one for each experimental condition (Repetition X Primeability; see color key at bottom). Each scatterplot shows 4 best-fit lines, one for each condition’s points separately. If variability along the y axis is due only to the corresponding nuisance factor, then all points should lie along a single best fit line with increasing slope. However, even if there is a strong effect of the nuisance factor, significant shifts along the y axis can indicate a real, residual effect of condition. The results for the ACC ROI in the right panels are consistent with all differences in similarity being due to the nuisance factors (note the highly similar best-fit lines, all with positive slope). Results for the Left Frontal ROI are less definitive, with a possible vertical shift in the position of the Strong Primeable, NEW fit line. However, it is also possible that the best-fit lines are simply noisy. The loss of findings for all other conditions in other ROIs suggests that SNR-level is poor enough that these results are, at best, indeterminate. (B) As anticipated, the average beta coefficient is related to estimated SNR, with beta values closer to zero associated with the lowest SNRs. The scatterplot shows all beta/SNR datapoints across all participants and ROIs, with significance testing by permutation (shuffling which beta goes with which SNR measure; 20,000 iterations). (C) The estimated SNRs from the current experiment lie in the range of 0.5 to 1.5, with a mean of approximately 1 (indicating equal levels of signal and noise). Given the likely presence of noise in the numerator of the calculations (see STAR methods), it is important to emphasize that these estimates are upper-bound estimates, which could mean the SNRs are actually poorer. In the plot, the estimated SNRs for the current MVPA data are shown relative to the theoretical curve of SNR-related attenuation of Pearson correlation coefficients (expressed as a fraction of the true correlation, e.g. 0.5; SNR on the x-axis actually depends on the individual SNRs of both patterns being correlated – see equation -- but is assumed to be the same for the two patterns here for simplicity). At SNRs of 1.0, small changes in SNR yield large differences in correlation (ranging from 0.2 to 0.6 of the true value when changing from SNRs of .5 to 1.5). At weaker values of SNR, this slope is even steeper. Taken together, it is possible that low beta weights lead to low SNR, which leads to low pattern similarity measures, potentially explaining the full pattern of raw similarity measures in Figure 8A. The level of SNR in single-trial fMRI responses may simply be too low to evaluate the Sharpening model in fMRI. Averaging responses across several repetitions, as is commonly done, is also inappropriate in this case, as testing the Sharpening model requires an estimation prior to any repetition. Related to Figure 8.

## References

Akaike, H. (1973). Information theory and an extension of the maximum likelihood principle. In Petrov, B.N. & Csáki, F. (eds.) 2^nd^ International Symposium on Information Theory, Tsahkadson, Armenia, USSR, September 2-8, 1971, Budapest: Akadémiai Kiadó, pp. 267–81.

Alink, A., Abdulrahman, H., & Henson, R.N. (2018). Forward models demonstrate that repetition suppression is best modelled by local neural scaling. Nat Commun 9, 3854. doi:10.1038/s41467-018-05957-0.

Atallah, H.E., Lopez-Paniagua, D., Rudy, J.W., & O’Reilly, R.C. (2007). Separate neural substrates for skill learning and performance in the ventral and dorsal striatum. Nat Neurosci 10, 126–31.

Bainbridge, W.A. (2017). The memorability of people: Intrinsic memorability across transformations of a person’s face. J Exp Psychol Learn Mem Cogn 43, 706–16.

Bainbridge, W.A. (2020). The resiliency of image memorability: A predictor of memory separate from attention and priming. Neuropsychologia 141:107408. doi:10.1016/j.neuropsychologia.2020.107408.

Bainbridge, W.A., & Rissman, J. (2018). Dissociating neural markers of stimulus memorability and subjective recognition during episodic retrieval. Sci Rep 8:8679. doi:10.1038/s41598-018-26467-5.

Balota, D.A., Yap, M.J., Corsese, M.J., Hutchison, K.A., Kessler, B., Loftis, B., Neely, J.H., Nelson, D.L., Simpson, G.B., and Trieman, R. (2007). The English lexicon project. Behav Res Meth 39, 445–459.

Bandettini, P.A. & Cox, R.W. (2000). Event-related fMRI contrast when using constant interstimulus interval: Theory and experiment. Magn Reson Med 43, 540–8.

Bartram, D.J. (1973). The effects of familiarity and practice on naming pictures of objects. Mem Cognit 1, 101–5.

Behzadi, Y., Restom, K., Liau, J., & Liu, T.T. (2007). A component based noise correction method (CompCor) for BOLD and perfusion based fMRI. NeuroImage 37, 90–101.

Berman, R.A., Gotts, S.J., McAdams, H.M., Greenstein, D., Lalonde, F., Clasen, L., et al. (2016). Disrupted sensorimotor and social–cognitive networks underlie symptoms in childhood-onset schizophrenia. Brain 139: 276–91.

Birn, R.M., Smith, M.A., Jones, T.B., & Bandettini, P.A. (2008). The respiration response function: the temporal dynamics of fMRI signal fluctuations related to changes in respiration. Neuroimage 40, 644–54.

Boettiger, C.A. & D’Esposito, M. (2005). Frontal networks for learning and executing arbitrary stimulus-response associations. J Neurosci 25, 2723–32.

Brunet, N.M, Bosman, C.A., Vinck, M., Roberts, M., Oostenveld, R., Desimone, R., De Weerd, P., & Fries, P. (2014). Stimulus repetition modulates gamma-band synchronization in primate visual cortex. Proc Natl Acad Sci U S A 111, 3626–31.

Bunzeck, N., Schütze, H., & Düzel, E. (2006). Category-specific organization of prefrontal response-facilitation during priming. Neuropsychologia 44, 1765–76.

Cabeza, R., Ciaramelli, E. & Moscovitch, M. (2012). Cognitive contributions of the ventral parietal cortex: An integrative theoretical account. Trends Cogn Sci 16, 338–52.

Cabeza, R., Mazuz, Y.S., Stokes, J., Kragel, J.E., Woldorff, M.G., Ciaramelli, E., Olson, I.R. & Moscovitch, M. (2011). Overlapping parietal activity in memory and perception: Evidence for the attention to memory model. J Cogn Neurosci 23, 3209–17.

Cave, C.B. (1997). Very long-lasting priming in picture naming. Psychol Sci 8, 322–5.

Cave, C.B., & Squire, L.R. (1992). Intact and long-lasting repetition priming in amnesia. J Exp Psychol Learn Mem Cogn 18, 509–20.

Chen, G., Glen, D.R., Saad, Z.S., Hamilton, J.P., Thomason, M.E., Gotlib, I.H., & Cox, R.W. (2011). Vector autoregression, structural equation modeling, and their synthesis in neuroimaging data analysis. Comput Biol Med 41, 1142–55.

Chen, G., Saad, Z.S., Britton, J.C., Pine, D.S., & Cox, R.W. (2013). Linear mixed-effects modeling approach to FMRI group analysis. Neuroimage 73, 176–90.

Cole, M.W., Pathak, S., Schneider, W. (2010). Identifying the brain’s most globally connected regions. NeuroImage 49, 3132–48.

Corbetta, M. & Shulman, G.L. (2011). Spatial neglect and attention networks. Annu Rev Neurosci 34, 569–99.

Cox, R.W. (1996). AFNI: software for analysis and visualization of functional magnetic resonance neuroimages. Comput Biomed Res 29, 162–73.

Cox, R.W., Chen, G., Glen, D.R., Reynolds, R.C. & Taylor, P.A. (2017). FMRI clustering in AFNI: False-positive rates redux. Brain Connect 7, 152–71.

De Baene, W., & Vogels, R. (2010). Effects of adaptation on the stimulus selectivity of macaque inferior temporal spiking activity and local field potentials. Cereb Cortex 20, 2145–65.

Desimone, R. (1996). Neural mechanisms for visual memory and their role in attention. Proc Natl Acad Sci U S A 93, 13494–9.

Dobbins, I.G., Schnyer, D.M., Verfaellie, M., & Schacter, D.L. (2004). Cortical activity reductions during repetition priming can result from rapid response learning. Nature 428, 316–9.

Eklund, A., Nichols, T., & Knutsson, H. (2016). Cluster failure: Why fMRI inferences for spatial extent have inflated false-positive rates. Proc Natl Acad Sci U S A 113, 7900–5.

Engell, A.D. & McCarthy, G. (2014). Repetition suppression of face-selective evoked and induced EEG recorded from the human cortex. Hum Brain Mapp 35, 4155–62.

Fischl, B., Salat, D.H., Busa, E., Albert, M., Dieterich, M., Haselgrove, C., van der Kouwe, A., Killiany, R., et al. (2002). Whole brain segmentation: automated labeling of neuroanatomical structures in the human brain. Neuron 33, 341–55.

Francis, W.S. (2014). Repetition priming in picture naming: sustained learning through the speeding of multiple processes. Psychol Bull Rev 21, 1301–8.

Friston, K.J. (1994). Functional and effective connectivity in neuroimaging: a synthesis. Hum Brain Mapp 2, 56–78.

Friston, K.J. (2005). A theory of cortical responses. Philos Trans R Soc Lond B Biol Sci 360, 815–36.

Friston, K.J. (2011). Functional and effective connectivity: a review. Brain Connect 1, 13–36.

Friston, K.J., & Kiebel, S.J. (2009). Predictive coding under the free-energy principle. Philos Trans R Soc Lond B Biol Sci 364, 1211–21.

Genovese, C., Lazar, N.A., & Nichols, T. (2002). Thesholding of statistical maps in functional neuroimaging using the false discovery rate. Neuroimage 15, 870–8.

Ghuman, A.S., Bar, M., Dobbins, I.G., & Schnyer, D.M. (2008). The effects of priming on frontal-temporal communication. Proc Natl Acad Sci U S A 105, 8405–9.

Gilbert, J.R., Gotts, S.J., Carver, F.W., & Martin, A. (2010). Object repetition leads to local increases in the temporal coordination of neural responses. Front Hum Neurosci 4: 30. doi:10.3389/fnhum.2010.00030.

Gilmore, A.W., Kalinowski, S.E., Milleville, S.C., Gotts, S.J., & Martin, A. (2019). Identifying task-general effects of stimulus familiarity in the parietal memory network. Neuropsychologia 124, 31–43.

Glover, G.H., Li, T.Q., Ress, D. (2000). Image-based method for retrospective correction of physiological motion effects in fMRI: RETROICOR. Magn Reson Med 44, 162–7.

Gotts, S.J. (2003). Mechanisms Underlying Enhanced Processing Efficiency in Neural Systems. Pittsburgh, PA: Carnegie Mellon University.

Gotts, S.J. (2016). Incremental learning of perceptual and conceptual representations and the puzzle of neural repetition suppression. Psychon Bull Rev 23, 1055–71.

Gotts S.J., Chow, C.C., & Martin, A. (2012). Repetition priming and repetition suppression: A case for enhanced efficiency through neural synchronization. Cogn Neurosci 3, 227–37.

Gotts, S.J., Gilmore, A.W., & Martin, A. (2020). Brain networks, dimensionality, and global signal averaging in resting-state fMRI: Hierarchical network structure results in low-dimensional spatiotemporal dynamics. Neuroimage 205:116289. doi:10.1016/j.neuroimage.2019.116289.

Gotts, S.J., Milleville, S.C., & Martin, A. (2015). Object identification leads to a conceptual broadening of object representations in lateral prefrontal cortex. Neuropsychologia 76, 62–78.

Gotts, S.J., Saad, Z.S., Jo, H.J., Wallace, G.L., Cox, R.W., & Martin, A. (2013). The perils of global signal regression for group comparisons: a case study of Autism Spectrum Disorders. Front Hum Neurosci Jul 12;7:356. doi:10.3389/fnhum.2013.00356.

Gotts, S.J., Simmons, W.K., Milbury, L.A., Wallace, G.L., Cox, R.W., & Martin, A. (2012). Fractionation of social brain circuits in autism spectrum disorders. Brain 135, 2711–25.

Graf, P., Squire, L. R., & Mandler, G. (1984). The information that amnesic patients do not forget. J Exp Psychol Learn Mem Cogn 10, 164–78.

Grill-Spector, K., Henson, R.N., & Martin, A. (2006). Repetition and the brain: neural models of stimulus-specific effects. Trends Cogn Sci 10, 14–23.

Haxby, J. V., Gobbini, M. I., Furey, M. L., Ishai, A., Schouten, J. L., & Pietrini, P. (2001). Distributed and overlapping representations of faces and objects in ventral temporal cortex. Science 293, 2425–30.

Henson, R.N. (2003). Neuroimaging studies of priming. Prog Neurobiol 70, 53–81.

Henson, R.N., Eckstein, D., Waszak, F., Frings, C. & Horner, A.J. (2014). Stimulus-response bindings in priming. Trends Cogn Sci 18, 376–84.

Henson, R.N., Horner, A.J., Greve, A., Cooper, E., Gregori, M., Simons, J.S., et al. (2017). No effect of hippocampal lesions on stimulus-response bindings. Neuropsychologia 103, 106–14.

Horner, A.J. (2012). Focusing on frontal cortex. Cogn Neurosci 3, 246–7.

Horner, A.J., & Henson, R.N. (2008). Priming, response learning and repetition suppression. Neuropsychologia 46, 1979–91.

Horner, A.J., & Henson, R.N. (2012). Incongruent abstract stimulus-response bindings result in response interference: fMRI and EEG evidence from visual object classification priming. J Cogn Neurosci 24, 760–73.

James, T. W., & Gauthier, I. (2006). Repetition-induced changes in BOLD response reflect accumulation of neural activity. Hum Brain Mapp 27, 37–45.

James, T.W., Humphreys, G.K., Gati, J.S., Menon, R.S., & Goodale, M.A. (2000). The effects of visual object priming on brain activation before and after recognition. Curr Biol 10, 1017–24.

Jasmin, K., Gotts, S.J., Xu, Y., Liu, S., Riddell, C.D., Ingeholm, J.E., Kenworthy, L., Wallace, G.L., Braun, A.R., & Martin, A. (2019). Overt social interaction and resting state in young adult males with autism: core and contextual neural features. Brain 142, 808–822.

Jo, H.J., Saad, Z.S., Simmons, W.K., Milbury, L.A. & Cox, R.W. (2010). Mapping sources of correlation in resting state FMRI, with artifact detection and removal. Neuroimage 52, 571–82.

Kaliukhovich, D.A., De Baene, W., & Vogels, R. (2013). Effect of adaptation on object representation accuracy in macaque inferior temporal cortex. J Cogn Neurosci 25, 777–89.

Kan, I.P., & Thompson-Schill, S.L. (2004). Effect of name agreement on prefrontal activity during overt and covert picture naming. Cogn Affect Behav Neurosci 4, 43–57.

Korzeniewska, A., Wang, Y., Benz, H.L., Fifer, M.S., Collard, M., Milsap, G., Cervenka, M.C., Martin, A., Gotts, S.J., & Crone, N.E. (2020). Changes in human brain dynamics during behavioral priming and repetition suppression. Prog Neurobiol Mar 18:101788. doi:10.1016/j.pneurobio.2020.101788.

Li, L., Miller, E.K., & Desimone, R. (1993). The representation of stimulus familiarity in anterior inferior temporal cortex. J Neurophysiol 69, 1918–29.

Maccotta, L., & Buckner, R.L. (2004). Evidence for neural effects of repetition that directly correlate with behavioral priming. J Cogn Neurosci 16, 1625–32.

McClelland, J.L., McNaughton, B.L., & O’Reilly, R.C. (1995). Why there are complementary learning systems in the hippocampus and neocortex: Insights from the successes and failures of connectionist models of learning and memory. Psychol Rev 102, 419–57.

McIntosh, A.R. & Gonzalez-Lima, F. (1994). Structural equation modeling and its application to network analysis in functional brain imaging. Hum Brain Mapp 2, 2–22.

McMahon, D.B., & Olson, C.R. (2007). Repetition suppression in monkey inferotemporal cortex: relation to behavioral priming. J Neurophysiol 97, 3532–43.

Meoded, A., Morrissette, A.R., Schanz, O., Gotts, S.J., & Floeter, M.K. (2015). Cerebro-cerebellar connectivity is increased in primary lateral sclerosis. Neuroimage Clin 7, 288–96.

Miller, E.K., Li, L., & Desimone, R. (1993). Activity of neurons in anterior inferior temporal cortex during a short-term memory task. J Neurosci 13, 1460–78.

Misaki, M., Kim, Y., Bandettini, P. A., & Kriegeskorte, N. (2010). Comparison of multivariate classifiers and response normalizations for pattern-information fMRI. NeuroImage 53, 103–18.

Mitchell, D.B. (2006). Nonconscious priming after 17 years: Invulnerable implicit memory? Psychol Sci 17, 925–9.

Newman, E.L., & Norman, K.A. (2010). Moderate excitation leads to weakening of perceptual representations. Cereb Cortex 20, 2760–70.

Norman, K.A., Newman, E.L., Detre, G.J., & Polyn, S.M. (2006). How inhibitory oscillations can train neural networks and punish competitors. Neural Comput 18, 1577–1610.

Norman, K.A., Polyn, S.M., Detre, G.J., & Haxby, J.V. (2006). Beyond mind reading: multi-voxel pattern analysis of fMRI data. Trends Cogn Sci 10, 424–30.

Op de Beeck, H.P., Torfs, K., & Wagemans, J. (2008). Perceived shape similarity among unfamiliar objects and the organization of the human object vision pathway. J Neurosci 28, 10111–23.

Packard, M.G. & Knowlton, B.J. (2002). Learning and memory functions of the basal ganglia. Annu Rev Neurosci 25, 563–93.

Power, J.D., Barnes, K.A., Snyder, A.Z., Schlaggar, B.L. & Petersen, S.E. (2012). Spurious but systematic correlations in functional connectivity MRI networks arise from subject motion. Neuroimage 59, 2142–54.

Price, L.R., Laird, A.R., Fox, P.T., & Ingham, R.J. (2009). Modeling dynamic functional neuroimaging data using structural equation modeling. Structural Equation Modeling 16, 147–62.

Race, E., Burke, K. & Verfaellie, M. (2019). Repetition priming in amnesia: Distinguishing associative learning at different levels of abstraction. Neuropsychologia 122, 98–104.

Race, E.A., Shanker, S., & Wagner, A.D. (2009). Neural priming in human frontal cortex: multiple forms of learning reduce demands on the prefrontal executive system. J Cogn Neurosci 21, 1766–81.

Ramirez, F.M. (2018). Orientation encoding and viewpoint invariance in face recognition: inferring neural properties from large-scale signals. Neuroscientist 24, 582–608.

Reid, A.T., Headley, D.B., Mill, R.D., Sanchez-Romero, R., Uddin, L.Q., Marinazzo, D., et al. (2019). Advancing functional connectivity research from association to causation. Nat Neurosci 22, 1751–60.

Saad, Z.S., Reynolds, R.C., Jo, H.J., Gotts, S.J., Chen, G., Martin, A., et al. (2013). Correcting brain-wide correlation differences in resting-state FMRI. Brain Connectivity 3, 339–52.

Salimpoor, V.N., Chang, C., & Menon, V. (2010). Neural basis of repetition priming during mathematical cognition: repetition suppression or repetition enhancement? J Cogn Neurosci 22, 790–805.

Salomon, R., Bleich-Cohen, M., Hahamy-Dubossarsky, A., Dinstien, I., Weizman, R., Poyurovsky, M., et al. (2011). Global functional connectivity deficits in schizophrenia depend on behavioral state. J Neurosci 31, 12972–81.

Sayres, R. & Grill-Spector, K. (2006). Object-selective cortex exhibits performance-independent repetition suppression. J Neurophysiol 95, 995–1007.

Scarborough, D.L., Cortese, C., & Scarborough, H.S. (1977). Frequency and repetition effects in lexical memory. J Exp Psychol Human 3, 1–17.

Schacter, D.L., & Buckner, R.L. (1998). Priming and the brain. Neuron 20, 185–95.

Seger, C.A. & Miller, E.K. (2010). Category learning in the brain. Annu Rev Neurosci 33, 203–19.

Smith, J.F., Pillai, A., Chen, K., & Horwitz, B. (2012). Effective connectivity modeling for fMRI: six issues and possible solutions using linear dynamical systems. Front Syst Neurosci Jan 18; 5:104. doi: 10.3389/fnsys.2011.00104.

Smith, R.E.W., Avery, J.A., Wallace, G.L., Kenworthy, L., Gotts, S.J. & Martin, A. (2019). Sex differences in resting-state functional connectivity of the cerebellum in Autism Spectrum Disorder. Front Hum Neurosci 13:104. doi:10.3389/fnhum.2019.00104.

Song, S., Gotts, S.J., Dayan, E., Cohen, L.G. (2015). Practice Structure Improves Unconscious Transitional Memories by Increasing Synchrony in a Premotor Network. J Cogn Neurosci 27, 1503–12.

Spearman, C. (1904). The proof and measurement of association between two things. Amer J Psychol 15, 72–101.

Squire, L.R. (1992). Memory and the hippocampus: a synthesis from findings with rats, monkeys, and humans. Psychol Rev 99, 195–231.

Stark, C.E., & McClelland, J.L. (2000). Repetition priming of word, pseudowords, and nonwords. J Exp Psychol Learn Mem Cogn 26, 945–72.

Steel, A., Song, S., Bageac, D., Knutson, K.M., Keisler, A., Saad, Z.S., et al. (2016). Shifts in connectivity during procedural learning after motor cortex stimulation: A combined transcranial magnetic stimulation/functional magnetic resonance imaging study. Cortex 74, 134–48.

Stoddard, J., Gotts, S.J., Brotman, M.A., Lever, S., Hsu, D., Zarate, C., et al. (2016). Aberrant intrinsic functional connectivity within and between corticostriatal and temporal–parietal networks in adults and youth with bipolar disorder. Psychol Med 46, 1509–22.

Suzuki, W.A. (2008). Associative learning signals in the brain. Prog Brain Res 169, 305–20.

Tong, F., Harrison, S.A., Dewey, J.A., & Kamitani, Y. (2012). Relationship between BOLD amplitude and pattern classification of orientation-selective activity in the human visual cortex. Neuroimage 63, 1212–22.

Tulving, E., & Schacter, D.L. (1990). Priming and human memory systems. Science 247, 301–6.

van Turennout, M., Bielamowicz, L., & Martin, A. (2003). Modulation of neural activity during object naming: Effects of time and practice. Cereb Cortex 13, 381–391.

van Turennout, M., Ellmore, T., & Martin, A. (2000). Long-lasting cortical plasticity in the object naming system. Nat Neurosci 3, 1329–34.

Warrington, E. K., & Weiskrantz, L. (1974). The effect of prior learning on subsequent retention in amnesic patients. Neuropsychologia 12, 419–28.

Watson, C.E., Gotts, S.J., Martin, A., & Buxbaum, L.J. (2019). Bilateral functional connectivity at rest predicts apraxic symptoms after left hemisphere stroke. Neuroimage Clin 21:101526. doi:10.1016/j.nicl.2018.08.033.

Weiner, K.S., Sayres, R., Vinberg, J., & Grill-Spector, K. (2010). fMRI-adaptation and category selectivity in human ventral temporal cortex: regional differences across time scales. J Neurophysiol 103, 3349–65.

Wig, G.S., Grafton, S.T., Demos, K.E., & Kelley, W.M. (2005). Reductions in neural activity underlie behavioral components of repetition priming. Nat Neurosci 8, 1228–33.

Wiggs, C.L., & Martin, A. (1998). Properties and mechanisms of perceptual priming. Curr Opin Neurobiol 8, 227–33.

Wiggs, C.L., Weisberg, J., & Martin, A. (2006). Repetition priming across the adult lifespan—the long and short of it. Neuropsychol Dev Cogn B Aging Neuropsychol Cogn 13, 308–25.

Xu, Y., Turk-Browne, N.B., & Chun, M.M. (2007). Dissociating task performance from fMRI repetition attenuation in ventral visual cortex. J Neurosci 27, 5981–5.

Yeo, B.T., Krienen, F.M., Sepulcre, J., Sabuncu, M.R., Lashkari, D., Hollinshead, M., Roffman, J.L., Smoller, J.W., et al. (2011). The organization of the human cerebral cortex estimated by intrinsic functional connectivity. J Neurophysiol 106, 1125–65.

Zachariou, V., Nikas, C.V., Safiullah, Z.N., Gotts, S.J., & Ungerleider, L.G. (2017). Spatial mechanisms within the dorsal visual pathway contribute to the configural processing of faces. Cereb Cortex 27, 4124–38.

