## Supplementary figures and images for "Enhanced inter-regional coupling of neural responses underlies long-term behavioral priming"

### Supplemental Figure 1

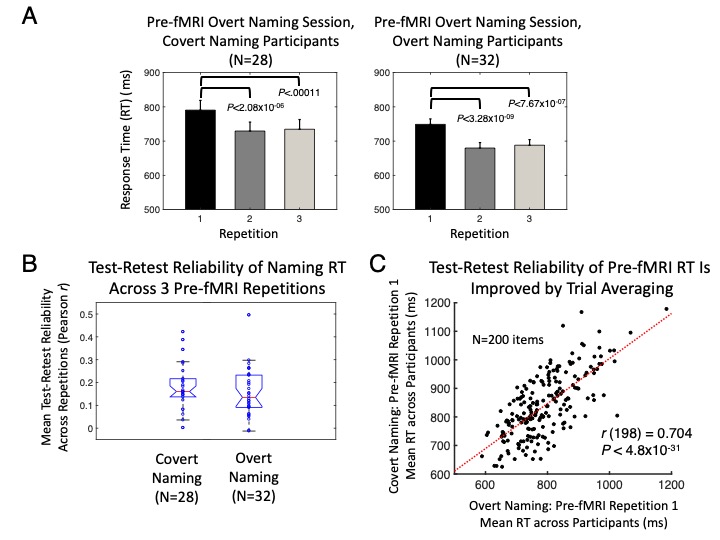

### Supplemental Figure 2

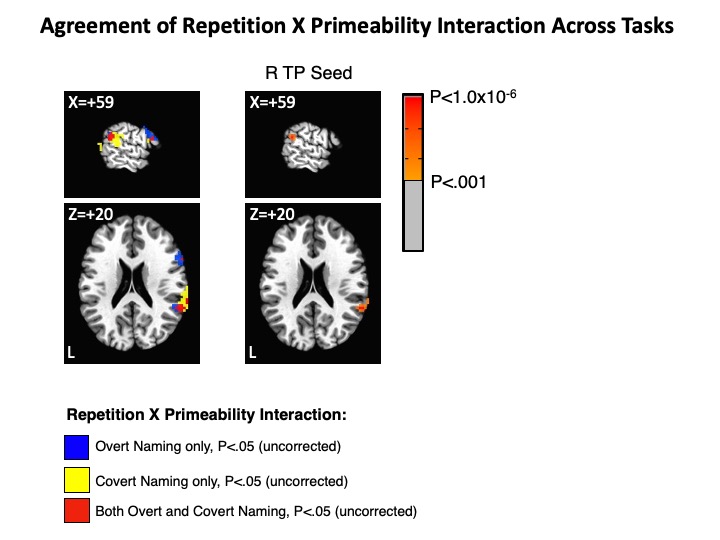

### Supplemental Figure 3

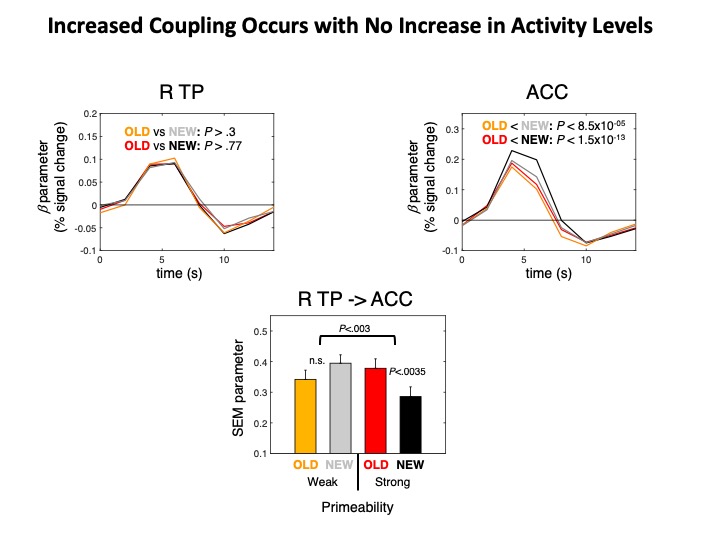

### Supplemental Figure 4

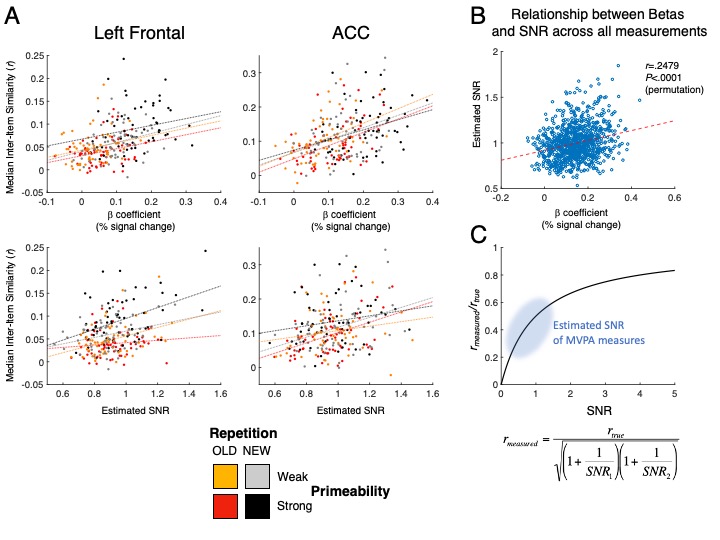
