## Supplemental Table 1 for "Enhanced inter-regional coupling of neural responses underlies long-term behavioral priming"

| | ROI Label | Peak Coordinate (Talairach-Tournoux) X | Y | Z | Spatial Extent RS effect (mm^3) | Spatial Extent FC effect (mm^3) | Overlap (mm^3) |
|---|---|---|---|---|---|---|---|
| 1 | Left Frontal | -40 | +5 | +26 | 11934 | | |
| 2 | Left Fusiform Gyrus | -40 | -49 | -10 | 5940 | | |
| 3 | Right Fusiform Gyrus* | +41 | -40 | -16 | 4455 | 1080 | 189 |
| 4 | Anterior Cingulate (ACC)* | +5 | +2 | +50 | 4752 | 891 | 324 |
| 5 | Right STG | +44 | -37 | +11 | | 918 | |
| 6 | Right Putamen | +29 | -16 | +8 | | 837 | |
| 7 | Right Temporoparietal | +59 | -43 | +20 | | 756 | |

* - Peak Coordinate used was from FC effect (in overlapping voxels with RS effect)
